# A Versatile Polypharmacology Platform Promotes Cytoprotection and Viability of Human Pluripotent and Differentiated Cells

**DOI:** 10.1101/815761

**Authors:** Yu Chen, Carlos A. Tristan, Lu Chen, Vukasin M. Jovanovic, Claire Malley, Pei-Hsuan Chu, Seungmi Ryu, Tao Deng, Pinar Ormanoglu, Dingyin Tao, Yuhong Fang, Jaroslav Slamecka, Christopher A. LeClair, Sam Michael, Christopher P. Austin, Anton Simeonov, Ilyas Singeç

## Abstract

Clinical translation of human pluripotent stem cells (hPSCs) requires advanced strategies that ensure safe and robust long-term growth and functional differentiation. Pluripotent cells are capable of extensive self-renewal, yet remain highly sensitive to environmental perturbations *in vitro*, posing challenges to their therapeutic use. Here, we deployed innovative high-throughput screening strategies to identify a small molecule cocktail that dramatically improves viability of hPSCs and their differentiated progeny. We discovered that the combination of Chroman 1, Emricasan, Polyamines, and Trans-ISRIB (CEPT) enhanced cell survival of genetically stable hPSCs by simultaneously blocking several stress mechanisms that otherwise compromise cell structure and function. In proof-of-principle experiments we then demonstrated the strong improvements that CEPT provided for several key applications in stem cell research, including routine cell passaging, cryopreservation of pluripotent and differentiated cells, embryoid body and organoid formation, single-cell cloning, genome editing, and new iPSC line generation. Thus, CEPT represents a unique polypharmacology strategy for comprehensive cytoprotection, providing a new rationale for efficient and safe utilization of hPSCs. Conferring cell fitness by multi-target drug combinations may become a common approach in cryobiology, drug development, and regenerative medicine.

## Introduction

Pluripotency is a remarkable cellular state that allows unlimited self-renewal and differentiation into all cell types of the human body. The isolation of the first hESC lines and the advent of the induced pluripotent stem cell (iPSC) technology have created novel paradigms for drug discovery, disease modeling, and cellular therapies^1–3^. However, to fully harness the therapeutic potential of iPSCs, it is critical to establish well-controlled, safe, and efficient strategies for cell line generation, cell expansion, directed differentiation into multiple phenotypes, and large-scale cell production (e.g. cryopreservation of master cell banks under good manufacturing practice conditions, “off-the-shelf” cellular products). Although significant progress has been made in characterizing the molecular and cellular features of pluripotency^4^, culturing hPSCs remains highly variable and uncontrolled^5–10^. This is particularly evident when hPSCs are dissociated into single cells for cell expansion, cryopreservation or cloning experiments^11^. Poor survival of dissociated cells and low efficiency of single-cell cloning are also major obstacles for genome editing in iPSCs^12^.

A landmark paper reported that the Rho-associated coiled-coil forming protein serine/threonine kinase (ROCK) inhibitor Y-27632 improves survival of hESCs^13^. Subsequent studies identified that Y-27632 and other small molecules that act on the ROCK pathway (e.g. Blebbistatin, Fasudil, Pinacidil, Thiazovivin) improve cell survival by blocking cell contraction^14–18^, which is detrimental to dissociated hPSCs and can lead to cell death by apoptosis and anoikis^19^. Interestingly, ROCK pathway inhibition also promotes the isolation and culture of sensitive cancer stem cells^20, 21^. Other fields that benefit from using Y-27632 include reproductive biology^22^, cryobiology^23^, and transplantation medicine^24, 25^. However, despite ROCK pathway inhibition and overall advances in cell culture techniques over the last two decades, poor cell survival remains a major roadblock due to lack of advanced approaches and incomplete understanding of the molecular mechanisms of cellular stress response in hPSCs. Hence, improving the viability of hPSCs and other sensitive cell types has become a daunting challenge. We here report the discovery and functional validation of a four-component small molecule cocktail, which enables integrated cytoprotection and provides new mechanistic insights into the complexity of cellular stress and can be broadly utilized in basic and translational research.

## Results

### Chroman 1 is superior to Y-27632 and other ROCK pathway inhibitors

To discover novel compounds that can promote cell survival, we first determined conditions for hPSCs that enable CellTiter-Glo (CTG) cell viability screening in highly miniaturized 1536-well plates (Z’ factor = 0.77; Fig. 1a and Supplementary Figs. 1, 2). Human ESCs (WA09) were plated onto vitronectin (VTN-N)-coated plates (500 cells/well) in chemically defined E8 medium and quantitative high-throughput screening (qHTS) was performed using diverse small molecule libraries^26, 27^. We screened a total of 15,333 compounds, and each compound was tested at 7 to 11 different concentrations to generate full dose-response curves^28^, a strategy not typically used in stem cell-based high-throughput screening studies. This initial screen yielded 113 hits and the primary targets of these hits were mainly protein kinases and proteases (Supplementary Figs. 3, 4 and Supplementary Table 1). Cluster analysis revealed that many active kinase inhibitors shared similar chemical structure with well-known ROCK1/2 inhibitors (Fig. 1b). Among these hits, 20 compounds were indeed known inhibitors of ROCK1/2 (e.g. Thiazovivin, Fasudil, GSK429286A). However, Chroman 1^29^ emerged as the most potent ROCK inhibitor (Fig. 1c) generating similar CTG readings as 10 μM Y-27632 when used at only 50 nM. To our knowledge, Chroman 1 has not been investigated in the context of stem cells so far. *In vitro* kinase assays confirmed that Chroman 1 was more potent against ROCK1 (IC_50_ = 52 pM) and ROCK2 (IC_50_ = 1 pM) than Y-27632 (ROCK1 IC_50_ = 71 nM and ROCK2 IC_50_ = 46 nM) (Fig. 1d,e). Next, using HotSpot kinase inhibitor profiling of 369 human kinases that cover all major human protein kinase families ^30^, we found that 10 μM Y-27632 inhibited not only its primary targets ROCK1/2 but also various off-targets such as PKCη (PRKCH), PKCε (PRKCE), PKCδ (PRKCD), PKN1, PKN2 and PRKX, to below 10% of their normal activity. In contrast, 50 nM Chroman 1 only inhibited ROCK1/2 to such levels while achieving efficient hPSC survival (Supplementary Figs. 5 and 6). Moreover, when hPSCs were dissociated and treated with both inhibitors for 24 h, significantly fewer dead cells (Figs. 1f,g, and 2b,c) and higher numbers of live cells (Fig. 2d) were detected in the presence of Chroman 1. Collectively, Chroman 1 was more potent and more selective against ROCK1/2 than Y-27632 and also improved cell survival by ∼25% (Fig. 2d). Based on these findings, we recommend replacing the widely used Y-27632 with Chroman 1.

**Figure 1.**
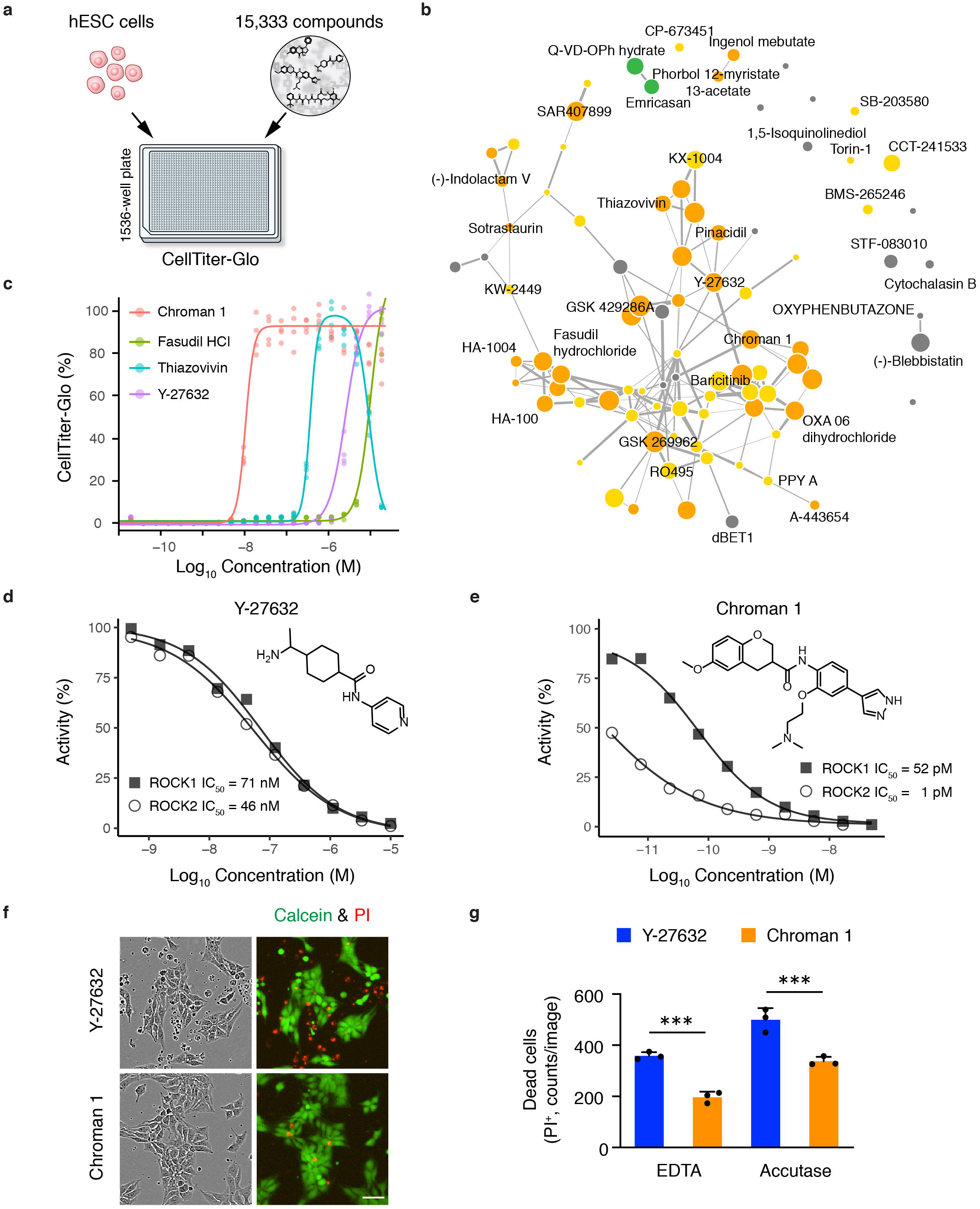
Quantitative HTS identifies Chroman 1 as a superior ROCK inhibitor. **a**, Cell survival assay in 1536-well format for qHTS. Cell viability was assessed after compound exposure for 24 h using CellTiter-Glo (CTG) to measure the cellular ATP levels of live cells. **b**, Chemical structure similarity analysis of active compounds. Point size correlates with maximum survival achieved by the compound at all tested concentrations. Two compounds are connected in the similarity network when their chemical structures are similar (Tanimoto coefficient > 0.17). Color indicates the primary target (orange = AGC protein kinases; yellow = other protein kinases, green = Caspases, gray = other targets). **c**, Dose-response curves of selected ROCK inhibitors, including Chroman 1, Fasudil, Thiazovivin and Y-27632 (n = 4 wells). CTG readings were normalized to the average number obtained with 10 μM Y-27632 (control). Note that Thiazovivin shows toxic effects at higher concentrations. **d, e,** Potency of Y-27632 and Chroman 1 against their primary targets ROCK1 and ROCK2 as determined by the HotSpot kinase assay (see also Supplementary Figs. 5 and 6). **f, g,** Improved survival of hESCs (WA09) with Chroman 1 treatment. Cells were dissociated with EDTA or Accutase and plated on vitronectin (VTN-N) in E8 medium at 100,000 cells/cm^2^. Phase contrast and fluorescence images were taken 12 h after plating. Live and dead cells were stained with calcein green AM and propidium iodide (PI), respectively. Data represent mean ± s.d., n = 3 wells for each group, and 36 fields of view were analyzed for each well. \*\*\**P* = 0.0004, one-way ANOVA. Scale bar in (**f**) 100 μm.

**Figure 2.**
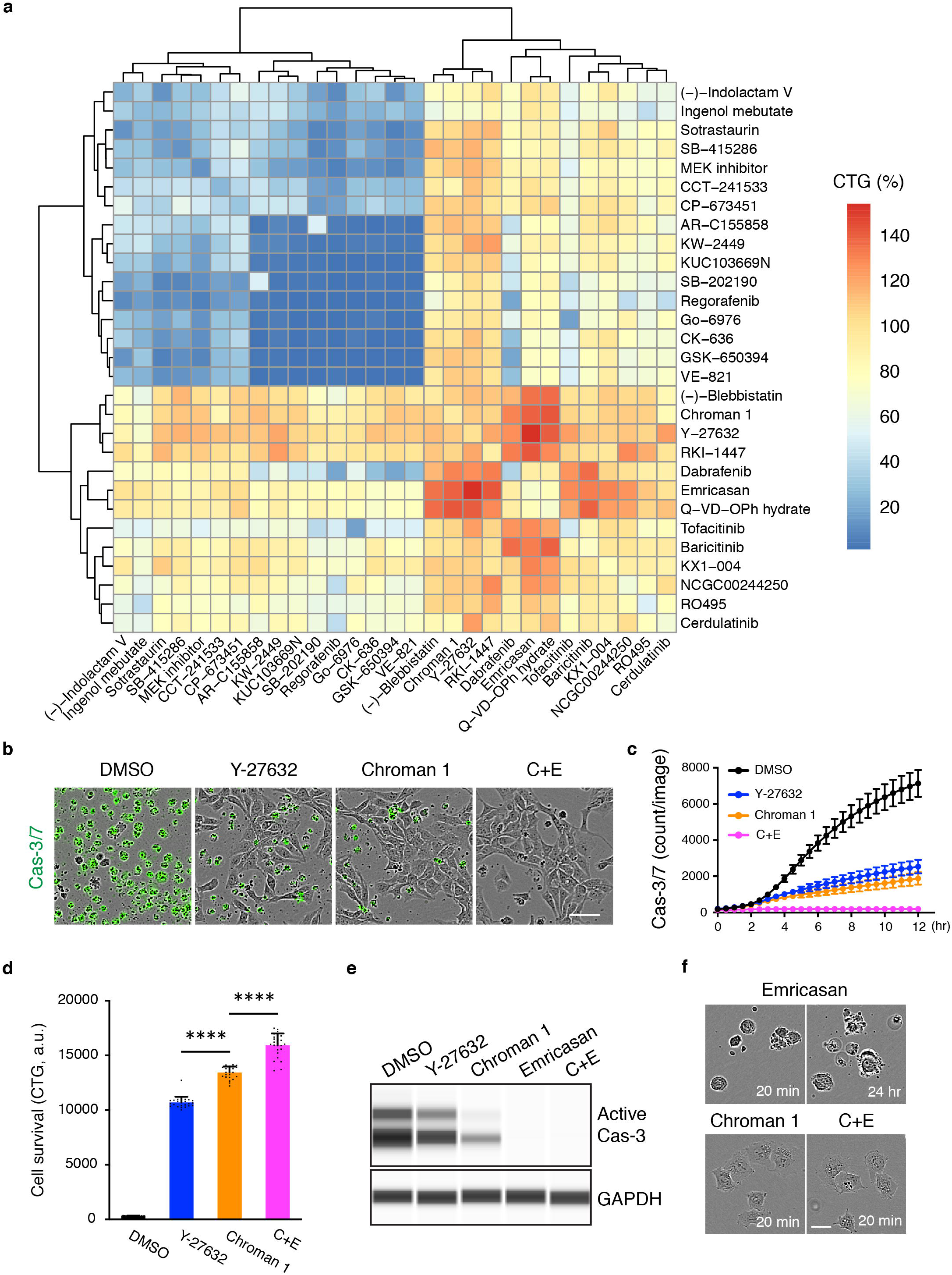
Combinatorial matrix screen identifies compounds with synergistic activities. **a**, Heatmap summarizing the maximum CTG readings of all-versus-all dose matrices for 29 compounds (812 drug-drug combinations tested in total; see also Supplementary Fig. 7 for examples showing 10 x 10 checkerboard dose matrices). CTG readings were normalized to the value obtained with 10 μM Y-27632 representing 100% (control). **b-d,** Combination of Chroman 1 with Emricasan (C+E) improves cell survival and reduces the number of apoptotic cells. hESCs (WA09) cells were dissociated with Accutase and plated on VTN-N in E8 medium (100,000 cells/cm^2^). Caspase-3/7 green detection reagent was used to monitor caspase activation (**b** and **c**) and CTG was used to quantify viable cells 24 h post-seeding (**d**). Data represent mean ± s.d. (**c**, n = 25 fields of view for each group; **d**, n = 24 wells for each group), \*\*\*\**P* < 0.0001, one-way ANOVA. Scale bar in **b**: 50 μm. **e**, Western blot analysis of Caspase-3 activation induced by single-cell dissociation. Cells were dissociated with Accutase and plated on VTN-N in the presence of indicated compounds. Cell lysates were collected after compound treatment for 2 h, and Caspase-3 activation (indicated by the cleaved version of Caspase-3) was measured. **f**, Live-cell imaging showing differences in cell behavior based on treatment with Emricasan, Chroman 1 or C+E. Cells were dissociated with Accutase and plated on VTN-N in the presence of indicated compounds. Cell blebbing continued in the presence of Emricasan and cells failed to attach to coated plates, and eventually underwent cell death. In contrast, hESCs readily attached within 20 min when treated with either Chroman 1 or C+E. Scale bar, 25 μm.

### Chroman 1 and Emricasan synergize to enhance cell survival

Next, we adopted a combinatorial matrix screening strategy^27^ to test whether synergistic drug combinations can further improve cell survival. Considering the diverse mode of action (MOA) of identified hits, we prioritized 29 compounds for matrix screening (Fig. 2a). This experiment generated 812 dose combination matrices (Supplementary Fig. 7 and https://ipsc.ncats.nih.gov/screening/). Using the Gaddum non-interacting model^27^, we identified several drug combinations with synergistic effects on cell survival (Fig. 2a). Generally, combinations of ROCK inhibitors or compounds with similar MOA (Chroman 1, Y-27632, RKI-1447, Blebbistatin) together with pan-Caspase inhibitors (Emricasan, Q-VD-OPh) yielded the strongest synergy scores (Supplementary Fig. 7a,b). In contrast, combined use of compounds with similar MOA (e.g. Chroman 1 and Blebbistatin) were modestly additive (Supplementary Fig. 7c,d).

Focusing on the combination of Chroman 1 plus Emricasan (C+E), we used three independent assays to confirm that C+E was superior to single compound treatment with either Y-27632 or Chroman 1. We monitored apoptosis following cell dissociation using CellEvent Caspase-3/7 detection reagent, which labels cells upon Caspase-3/7 activation. Exposure to C+E greatly reduced the emergence of green fluorescence, indicating minimal cell death (Fig. 2b,c). Accordingly, C+E treatment resulted in the highest number of live cells (CTG assay) among all treatment groups (Fig. 2d; ∼48% improvement over Y-27632). Western blots showed that Y-27632 or Chroman 1 partially inhibited Caspase-3 activation in dissociated cells (Fig. 2e), whereas C+E treatment completely blocked Caspase-3 activation (Fig. 2e). Emricasan alone also efficiently blocked Caspase-3 activation (Fig. 2e); however, without ROCK inhibition ongoing cell contractions eventually led to cell death (Fig. 2f and Supplementary Movie 1).

To demonstrate that transient exposure to Emricasan was safe and did not permanently alter apoptosis in hPSCs, we monitored the recovery of Caspase-3 activation after Emricasan washout. We found that active Caspase-3 returned to normal levels 24-48 h after removing Emricasan, while total Caspase-3 levels remained unaffected (Supplementary Fig. 8). Furthermore, to ensure that the use of Chroman 1 or C+E was safe, we cultured hESCs and iPSCs for over 40 passages and treated cells at every passage for 24 h with either Chroman 1 or C+E. These cells maintained normal karyotypes, high expression levels of pluripotency markers OCT4 and NANOG, and showed multi-lineage differentiation potential in monolayer and EB cultures (Supplementary Fig. 9).

### High-throughput screening at ultra-low cell density identifies the four-factor cocktail CEPT

Cell density and cell-cell contact are key determinants for the survival of hPSCs^9, 31^. The use of Chroman 1 alone or C+E significantly improved hPSC survival over Y-27632 during routine cell passaging (∼25% and ∼48%, respectively), however, advancement was only modest under extreme conditions such as very low cell density (25 cells/cm^2^; compare Y-27632 and C+E in Fig. 3e,f). This prompted us to develop a new combination screening assay to search for additional compounds that, when applied together with C+E, can further enhance cell survival. In this assay, we seeded only 10 cells into each well of 1536-well plates (500 cells/well was used in our original screen; Supplementary Figs. 1, 2) to model the stress associated with ultra-low cell densities (Fig. 3a and Supplementary Fig. 10). To our knowledge, high-throughput combination screening at ultra-low cell density has not been performed in the published literature. We used 3 replicate sets of plates to compensate for the variation that occurs when seeding such few cells for HTS. Using this new assay, a total of 7599 compounds was screened. After ranking all compounds by the median CTG values, we prioritized 316 hits that appeared to improve cell survival (Fig. 3b). Guided by dose-response curves (Fig. 3c), we selected 36 hits for further validation. Quantification of cell survival revealed that Trans-ISRIB had the most significant effect when administered together with C+E (Fig. 3d). Trans-ISRIB is a selective inhibitor of the integrated stress response (ISR) and protein kinase RNA-like endoplasmic reticulum kinase (PERK) signaling, thereby disinhibiting protein translation stalled by phosphorylated eIF2A^32–34^. Interestingly, other PERK inhibitors such as AMG PERK 44, GSK2606414, or GS2656157 did not show the same effect as Trans-ISRIB (Fig. 3b). This suggests that Trans-ISRIB has unique properties and can enhance survival of hPSCs when used in combination with C+E (CET; Fig. 3e,f). However, Trans-ISRIB did not improve cell survival when administered alone in our original single-agent screen (Supplementary Table 1), further underscoring the importance of polypharmacology and identification of synergistic drug combinations. Quantification confirmed the positive effect of CET on clone number and size (Fig. 3e,f). Moreover, by testing additional reagents and conditions (e.g. 5% low oxygen, Knockout Serum Replacement, Albumax, ascorbic acid, antioxidant supplement), we found that a mixture of Polyamines (40 ng/mL putrescine, 4.5 ng/mL spermidine, 8 ng/mL spermine) together with C+E (CEP) had beneficial effects on single-cell cloning as well (Fig. 3e,f). Polyamines are essential polycations that modulate a number of cellular functions such as transcription, translation, cell cycle, stress response^35^ and self-renewal of mouse ESCs^36^. Collectively, under ultra-low cell density conditions the combination of <u>C</u>hroman 1, <u>E</u>mricasan, <u>P</u>olyamines and <u>T</u>rans-ISRIB (CEPT) showed the most robust effects on cell survival and cloning efficiency (Fig. 3e-g).

**Figure 3.**
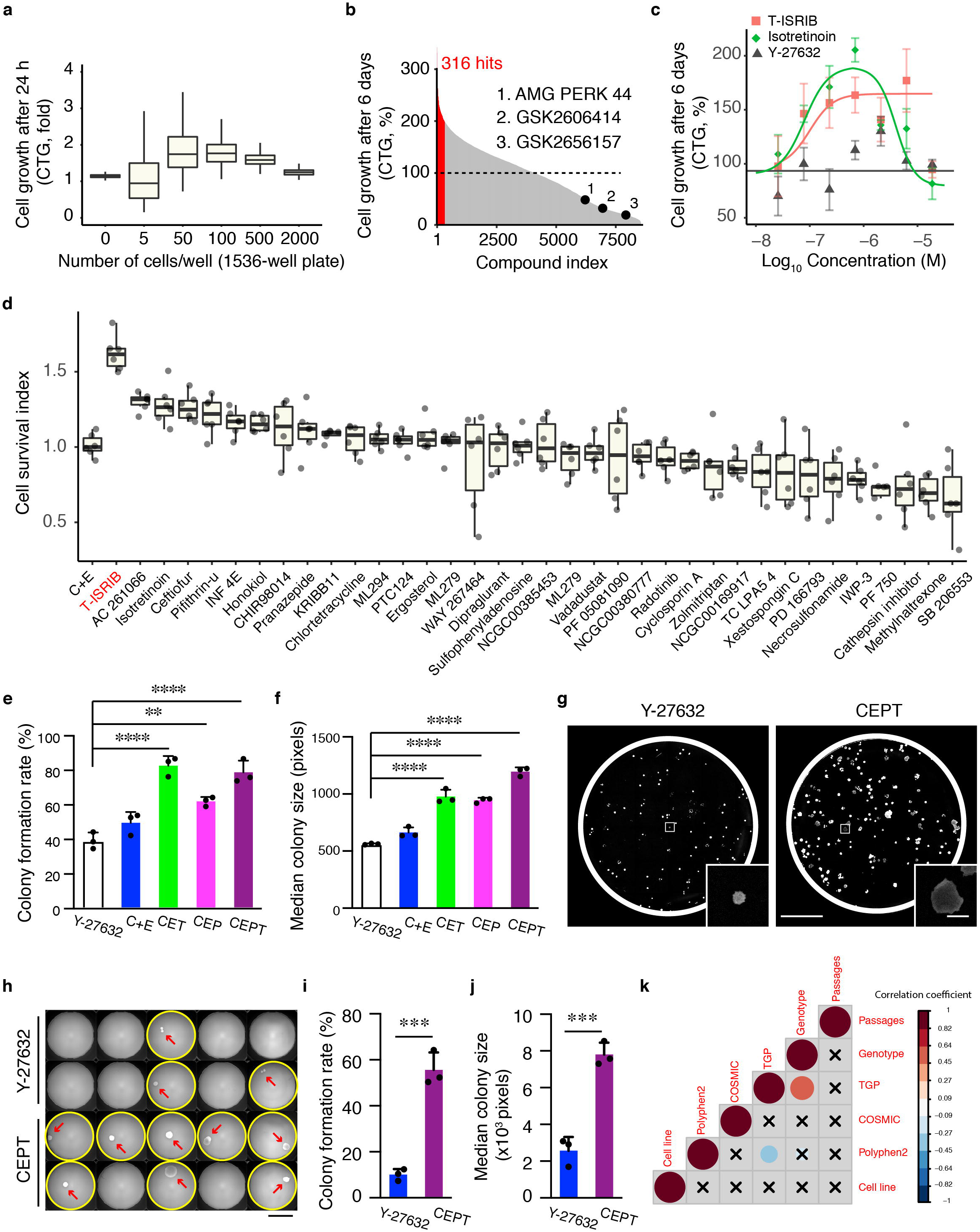
Combination of Chroman 1, Emricasan, Polyamines and Trans-ISRIB promote clonal growth and expansion of genetically stable hPSCs. **a**, Assay development for hESCs (WA09) in 1536-well format to model stress associated with ultra-low cell density on survival and growth. Fewer than 50 cells/well was determined as ultra-low cell density (see also Supplementary Fig. 10 for more details). **b**, Summary of qHTS performed at ultra-low cell seeding density (10 cells/well). All compounds were used in combination with C+E, tested at 10 μM in triplicates, and ranked based on their median CTG readings. Data were normalized to the average CTG reading obtained with C+E. 316 hits showed synergy with C+E (see Supplementary Table 2 for details). Note that the three indicated stress pathway inhibitors (PERK inhibitors) did not show synergy with C+E. **c**, Example dose-response curves of Trans-ISRIB, Isotretinoin, and Y-27632. Data represent median ± s.d. (n = 6 wells). **d**, Superiority of Trans-ISRIB in promoting growth at ultra-low cell density. hESCs (WA09) were plated on LN521 at 25 cells/cm^2^ in StemFlex medium with the indicated compounds in addition to C+E. After a 3-day incubation with compounds, cell confluency was quantified using calcein green AM to label live cells on day 6. Thirty six primary hits were followed up based on data presented in **b** and **c**. Data are presented as box plot (n = 6 wells for each group). Cell survival index represents cell confluency normalized to the C+E group. **e-g,** Systematic comparison of small molecule combinations and their effect on colony number and colony size of hESCs plated on VTN-N at 25 cells/cm^2^ in StemFlex medium. Quantification was performed 6 days after plating. Note that CET (Chroman 1, Emricasan, Trans-ISRIB), CEP (Chroman 1, Emricasan, Polyamines), and CEPT (Chroman 1, Emricasan, Polyamines, Trans-ISRIB) are superior to Y-27632 and C+E (Chroman 1 and Emricasan). Scale bar in (**g**), 8 mm; inset, 1 mm. In **e** and **f**, data are mean ± s.d. (n = 3 wells for each group), \*\*\*\**P* < 0.0001, \*\**P* = 0.0033, one-way ANOVA. **h,** Single cell cloning experiment in 96-well plates. Single cells were deposited using a cell sorter on VTN-N in StemFlex medium and treated with Y-27632 or CEPT for 3 days. Whole-well images were captured with calcein green AM to quantify clone numbers and size on day 9. Arrows show single clones in each well. Scale bar, 3.2 mm. **i, j**, Quantification of single cell cloning experiment. Data represent mean ± s.d. (three 96-well plates for each group), \*\*\**P* = 0.0006 in (**i)** and 0.0008 in (**j)**, two-tailed Student’s *t*-test. **k**, Pearson correlogram and significance of 95% confidence interval (X = not significant, color indicates direction of correlation). There was no correlation between cell line, passage number, or SNP genotype.

### Efficient and safe single-cell cloning and cell propagation by CEPT

To perform accurate single-cell cloning experiments, we first assessed how the mechanical stress of cell sorting may affect hPSC survival. Testing cell membrane integrity by using propidium iodide (PI) before and after passing cells through the sorter, we noted that ∼40% of the cells were immediately killed by the sorting process (Supplementary Fig. 11). Mechanical stress due to cell sorting is therefore a technical challenge for efficient cloning of hPSCs. This confounder was taken into account when single cell cloning efficiency of hPSCs was determined in 96-well plates following cell sorting, where CEPT treatment again showed strong improvement compared to Y-27632 (cloning efficiency 9% for Y27632 and 55% for CEPT; Fig. 3h-j). Eight clones generated with CEPT were randomly selected and established as clonal cell lines. After 4 passages, all clonal cell lines (8/8) were karyotypically normal (Supplementary Fig. 12) suggesting that single cell cloning using CEPT did not cause apparent genetic abnormalities. Next, whole exome sequencing (WES) was used to monitor hESCs and iPSCs treated with CEPT for 20 passages (CEPT treatment for 24 h at every passage). Both cell lines maintained normal features of pluripotency including self-renewal and multi-lineage differentiation (Supplementary Fig. 13). To perform more detailed genetic analysis on cells passaged with CEPT, we annotated variants for known cancer hotspots using various databases such as Cancer Hotspots^37^, Catalogue of Somatic Mutations in Cancer (COSMIC)^38^; hg38 RefSeq Gene^39^, dbSNP150^40^, The 1000 Genomes Project (TGP)^41^; The Exome Aggregation Consortium (ExAC)^42^, PolyPhen2^43^, and ClinVar^44^ (Fig. 3k and Supplementary Fig. 14). Recently reported dominant-negative mutations in the tumor suppressor gene TP53 ^10^ were absent in hESCs and iPSCs, before and after CEPT treatment (passage 0, 10, 20). The only detected exonic TP53 single nucleotide polymorphism (SNP), rs1042522, was already present in hESCs and iPSCs, did not change due to CEPT exposure during passaging (Supplementary Fig. 14a), and is likely to be benign (ClinVar). Genotypes of 19 other COSMIC variants that were exonic and nonsynonymous also did not change during passaging (Supplementary Fig. 14a). Similarly, there was no significant correlation between carrier status of single SNPs or insertions/deletions (indels), number of passages after CEPT treatment, cell line, predicted variant effect, or sample genotype (Fig. 3k and Supplementary Fig. 14b-d). These findings suggest that serial passaging with CEPT did not cause widespread genetic abnormalities or cancer-causing mutations in hESCs and iPSCs.

### Efficient genome editing of cloned hPSCs

Genome editing of hPSCs is cumbersome and time-consuming because clonal survival of hPSCs is inefficient, requiring additional strategies such as drug selection, genetic modifications, and laborious cell culture work to expand and identify precisely edited cells^12, 45–47^. Having established CEPT as a powerful tool for cell cloning (Fig. 3e-j), we sought to demonstrate its usefulness for genome editing using CRISPR/Cas9. As proof-of-principle, we followed a recently published protocol^45^ and used the same parental cell line and reagents in order to engineer a knock-in reporter cell line by introducing the sequence for GFP in front of exon 1 of the LMNB1 gene (Fig. 4a). After nucleofection of the donor vector together with the ribonuclear protein (RNP) complex of wild-type Cas9 protein and a synthetic sgRNA targeting the first exon of LMNB1, we obtained a mixture of GFP^+^ and GFP^-^ cells at day 3 (Fig. 4b). GFP^+^ cells displayed varying fluorescence intensities and labeling of the peri-nuclear region suggested successful tagging of GFP to the N-terminus of LMNB1 (Fig. 4b). To identify correctly edited cells, individual GFP^+^ cells were sorted into single wells of 96-well plates. Treatment with CEPT enabled robust cell survival and clone formation. On day 12, we randomly picked 8 colonies and established clonal lines, which again was facilitated by CEPT administration during clone picking and subsequent passaging. Droplet digital PCR (ddPCR) was used to quantify copy numbers of the GFP-tag sequence, the ampicillin resistance gene (AMP) as part of the vector backbone, and the two-copy genomic reference gene *RPP30*. Of these 8 clones, five clones showed mono-allelic integration, one clone indicated bi-allelic integration, and two clones had integrated the vector backbone containing AMP (Fig. 4c,d). Fluorescence intensity, junctional PCR, Sanger sequencing and Western blot analysis of clone #5 (mono-allelic) and #8 (bi-allelic) confirmed accurate editing and differential expression of GFP-LMNB1 (Fig. 4e-i). Moreover, genome-edited clones were karyotypically normal and capable of multi-lineage differentiation (Supplementary Fig. 15). While our strategy using CEPT enabled efficient and rapid clonal cell line establishment from accurately edited cells, less stringent conditions generating mosaic clones may require testing hundreds of putative clones^45^. Hence, CEPT should facilitate the streamlined production of genome-edited iPSC lines for basic and translational research.

**Figure 4.**
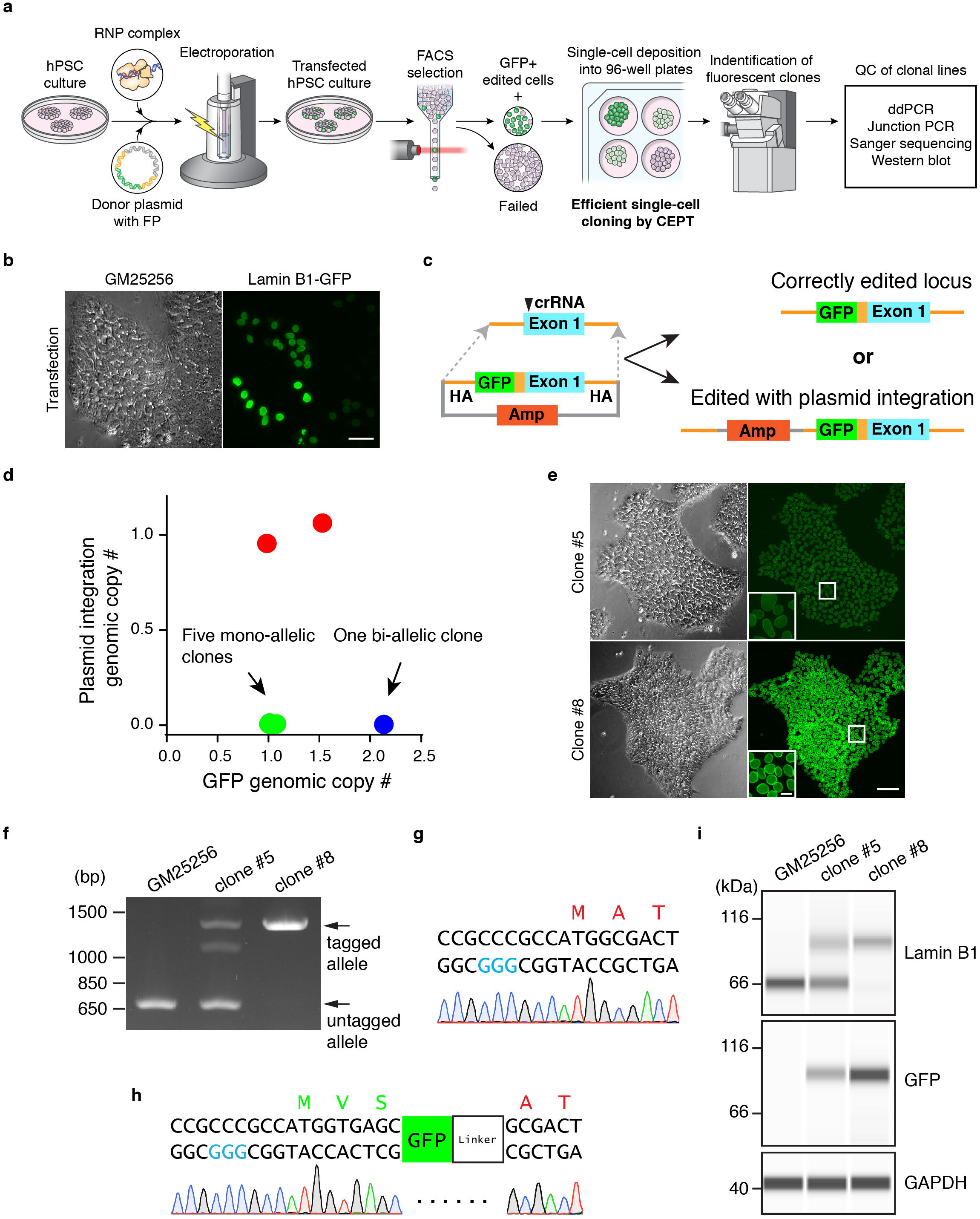
Efficient gene editing in cloned iPSCs. **a,** Workflow to generate genome-edited clonal cell lines from iPSCs. **b**, Detection of GFP^+^ cells 3 days after electroporation. Scale bar, 5 μm. **c,** Schematic of potential gene editing results with or without the integration of the plasmid backbone (AMP), both of which may lead to the expression and correct localization of GFP. **d,** Randomly picked GFP^+^ clones (n = 8) were analyzed for genomic copy numbers of GFP and the plasmid backbone (AMP) using ddPCR. One clone showed bi-allelic correct editing (blue circle), five clones showed mono-allelic correct editing (green circles), and two clones were edited with plasmid integration (red circles). **e,** Microscopic images of one of the mono-allelic correctly edited clones (clone #5) and the bi-allelic clone (clone #8). Note the stronger GFP signal intensity in the bi-allelic clone. Scale bar, 10 μm; inset, 2 μm. **f,** Junction PCR to detect the GFP insertion into the N-terminus of LMNB1 in clone #5 and #8. GM25256 represents the parental iPSC line. **g, h**, Sanger sequencing confirming edited and unedited alleles. **i**, Western blot analysis of LMNB1 expression in cell lines with mono- and bi-allelic modification and the parental iPSC line (GM25256).

### Optimizing EB and organoid formation by CEPT

Generation of three-dimensional EBs from hPSCs is a widely used assay to measure pluripotency and differentiation into the three germ layers^48, 49^. More recently, EBs have been used to produce organoids from hPSCs^50, 51^. Since large numbers of cells undergo cell death and introduce significant variability during EB and organoid formation, we asked if CEPT could be beneficial in this context. Human ESCs were dissociated and plated into ultra-low attachment (ULA) plates in the presence of DMSO, Y-27632, or CEPT. After 24 h, a striking difference was observed in that CEPT produced not only more numerous EBs but these cultures were also virtually devoid of cellular debris (Fig. 5a). Dramatic improvement in cell survival was further confirmed under more controlled conditions when single EBs were generated in individual wells. Exposure to CEPT for 24 h generated larger EBs (Fig. 5b,c), minimized cell death (Fig. 5d), and improved cell survival by 242% as compared to Y-27632 (Fig. 5e). Importantly, robust and continuous cell growth over 7 days was observed in EBs initiated with CEPT but not with Y-27632 (Fig. 5e). Next, we asked if improving EB quality due to minimal cell death might have functional consequences. Using 96-well plates, 20,000 cells were plated in chemically-defined E6 medium to form single EBs per well. At day 7, single EBs were measured for the expression of ectodermal (PAX6), mesodermal (Brachyury), and endodermal genes (SOX17). Notably, to enable the analysis of such low amounts of mRNA yielded by individual EBs, we developed an optimized quantitative PCR protocol (see Methods for details). Indeed, CEPT treatment more reliably generated EBs capable of differentiating into all three germ layers as compared to Y-27632 (Fig. 5f). Hence, the proper analysis of pluripotency using EBs can be improved by CEPT.

**Figure 5.**
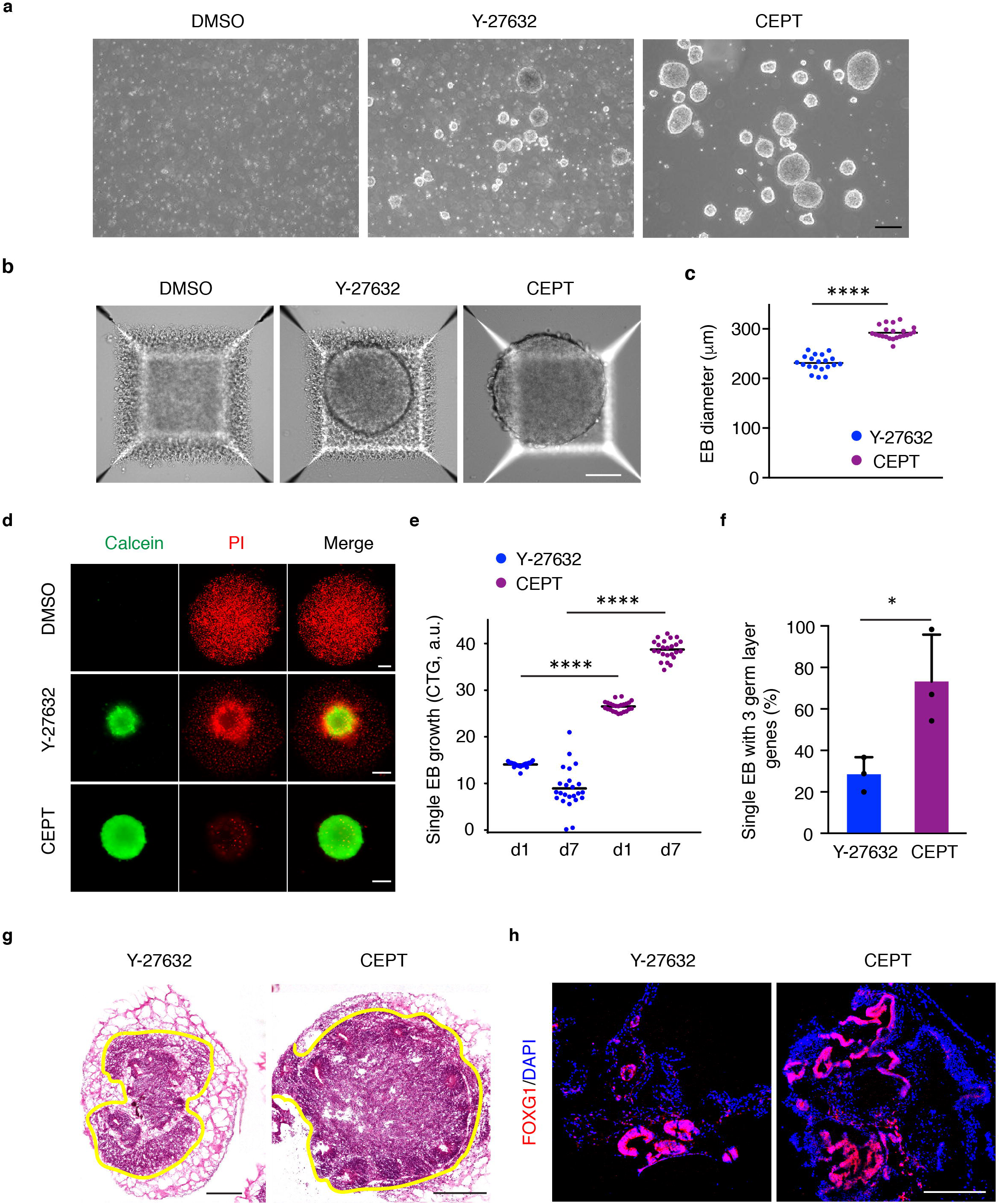
Optimized EB and organoid formation. **a,** EB formation in the presence of DMSO, Y-27632 and CEPT. Human ESCs (WA09) cells were dissociated with Accutase and plated into 6-well ULA plates in E6 medium. Representative phase-contrast images were taken at 24 h post-plating. Scale bar, 50 μm. **b,** To generate single EBs, hESCs were dissociated with Accutase and plating into AggreWell plates (5,000 cells/well). Images were taken 24 h post-plating. In the presence of DMSO the vast majority cells underwent cell death and EB formation was not observed. Treatment with Y-27632 supported EB formation but significant numbers of dead cells were detected surrounding the EB. Note that CEPT enables EB formation without apparent cell death. Scale bar, 100 μm. **c,** Quantification of the diameter of single EBs (24 h post-plating). Data are mean ± s.d. (n = 20 EBs for Y-27632 and n = 22 for CEPT), \*\*\*\**P* < 0.0001, two-tailed Student’s *t*-test. **d,** Single EB formation in 96-well ULA plates. Dissociated hESCs were plated into 96-well ULA plates at 2,000 cells/well. Live and dead cells were stained with calcein green AM and propidium iodide (PI) 24 h after cell seeding. Scale bars, 100 μm. **e,** Quantification of cell survival in single EBs at day 1 and day 7 by using the CTG 3D assay. Note the significant difference between Y-27632 and CEPT treatment at both timepoints. Data represent mean ± s.d. (n = 24 EBs for each group), \*\*\*\**P* < 0.0001, one-way ANOVA. **f,** CEPT supports differentiation of single EBs into the three germ layers. Individual EBs were cultured in E6 medium to allow for spontaneous differentiation and analyzed on day 7 for the expression of PAX6, SOX17, and Brachyury using an optimized quantitative RT-PCR protocol that enabled detection of low transcript levels in single EBs (see Materials and Methods section for details). Data represent mean ± s.d. (n = 3 experiments and in each experiment 24 EBs were analyzed for each group), \**P* = 0.0327, two-tailed Student’s *t*-test. **g, h,** Cerebral organoids were generated by using Y-27632 and CEPT for the first 24 h. At day 30, organoids were fixed, sectioned, processed for histology (hematoxylin and eosin stain) and immunohistochemistry for FOXG1. Representative images show that CEPT treatment resulted in larger organoids and more abundant FOXG1-expressing cells. Scale bars, 400 μm.

To investigate if organoid formation may also benefit from improved cell survival, we generated cerebral organoids following a published protocol^50^. Histological analysis using hematoxylin-eosin staining at day 30 showed a dramatic difference in neural tissue differentiation in cerebral organoids that were either initiated with Y-27632 or CEPT (Fig. 5g). Immunohistochemistry for the telencephalic marker FOXG1^50^ further confirmed that organoids generated by using CEPT produced larger amounts of neural tissue displaying complex morphologies (Fig. 5h). In summary, these findings underscore that optimizing cell survival during EB and organoid formation within the first 24 h has long-lasting consequences. Controlling cell survival and minimizing cell death as an inherent variable should help with developing new assays that may enable more faithful analysis of pluripotency and organoid-based disease modeling.

### A defined chemical foundation for regenerative medicine

The results described above suggested that CEPT might have broader applicability in regenerative medicine. Development of cellular products requires reliable protocols for biobanking of various phenotypes at different developmental stages for on-demand use in drug screening, toxicology, and cell therapies. Cryopreservation and thawing hPSCs has been notoriously problematic and inefficient^11, 52, 53^. Many scientific groups, including ours, have been using ROCK inhibitors as the only available option, although the amount of cell death remains substantial even in the presence of Y-27632. Remarkably, thawing cryopreserved hPSCs was dramatically improved after CEPT treatment for 24 h (∼300%) compared to Y-27632 (Fig. 6a-c). Next, we asked if CEPT may also have beneficial effects when thawing cryopreserved differentiated cells. Several iPSC-derived cell types including cardiomyocytes, hepatocytes, astrocytes and motor neurons showed significantly improved cell survival due to CEPT treatment (Fig. 6d). For instance, cardiomyocytes showed ∼36% improvement and motor neurons exhibited ∼63% improvement as compared to the DMSO control group. Notably, improved survival of cardiomyocytes and motor neurons had long-lasting consequences and was reflected by enhanced functional activity as measured by multi-electrode arrays for up to 14 days post-plating (Fig. 6e-g and Supplementary Fig. 16).

**Figure 6.**
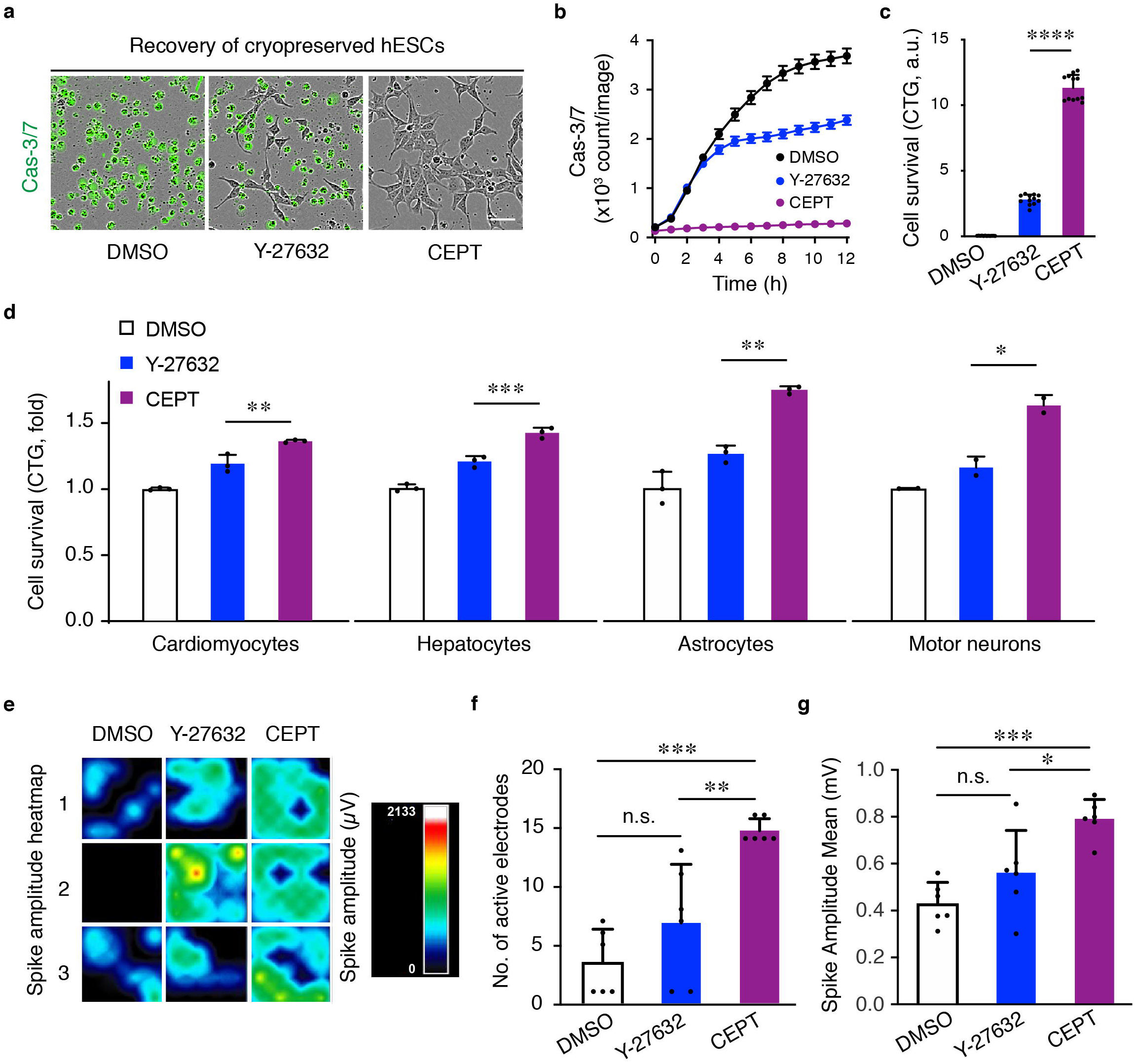
CEPT enables superior cryopreservation and improved function post-thawing. **a-c,** Cryopreserved hESCs (WA09) were thawed and plated in E8 medium in the presence of indicated compounds. Caspase-3/7 green detection reagent was used to monitor apoptosis over 12 h (**a** and **b**) and the CTG assay was used to quantify live cells 24 h post-thawing (**c**). Data represent mean ± s.d. (**b**, n = 25 fields of view for each group; **c**, n = 12 wells for each group), \*\*\*\**P* < 0.0001, one-way ANOVA. Scale bar in **a**, 50 μm. **d,** Frozen vials of iPSC-derived cardiomyocytes, hepatocytes, astrocytes, and motor neurons were thawed and treated with DMSO, Y-27632, and CEPT for 24 h. Cell survival was quantified using the CTG assay. Data are mean ± s.d. (n = 3 wells for each group of cardiomyocytes, hepatocytes and astrocytes, and n = 2 wells for motor neurons), \**P* < 0.05, \*\**P* < 0.005, \*\*\**P* < 0.001, one-way ANOVA. **e-g**, Electrophysiological characterization of iPSC-derived cardiomyocytes 5 days post-thawing using multi-electrode arrays. In **f**, data represent mean ± s.d. (n = 6 wells for each group), \*\**P* = 0.0029, \*\*\**P* = 0.0001, one-way ANOVA. In **g**, data represent mean ± s.d. (n = 6 wells for each group), \**P* = 0.0165, \*\*\**P* = 0.0005, one-way ANOVA.

Lastly, to determine whether CEPT can support new iPSC line generation, we used a Sendai virus-based method and the Yamanaka factors (OCT4, SOX2, C-MYC, KLF4) to reprogram skin fibroblasts^54^. Picking clones or colonies and establishing new iPSC lines is a critical step influenced by multiple technical and biological variables (e.g. donor age, proliferation rate, colony picking technique, culture conditions)^55^. When individual, similar-sized colonies (n = 30) of reprogrammed cells were picked and carefully transferred to single wells of 6-well plates (10 colonies per condition), CEPT treatment yielded significantly more cellular progeny than the DMSO and Y-27632 groups (Supplementary Fig. 17). Thus, CEPT should facilitate colony picking and iPSC line establishment as part of routine protocols and large-scale approaches^56^.

### Mechanistic insights into comprehensive cytoprotection

Previous reports suggested that hPSCs, as compared to differentiated cells, display an overall increased propensity for cell death following replication stress or DNA damage^57, 58^. Interestingly, the basal expression level of total Caspase-3 was higher in pluripotent cells and down-regulated during differentiation into ectoderm, mesoderm, and endoderm (Supplementary Fig. 18). Note that Emricasan is part of CEPT and inhibits Caspase-3 activation (Fig. 2 and Supplementary Fig. 8). Next, to gain insights into the cellular and molecular mechanisms that are initiated in hPSCs after cell dissociation, we performed a series of complementary experiments. Irregularly shaped nuclei of stressed cells (nuclear dysmorphia) can compromise genomic stability in normal and cancerous cells^59^. Using a lamin B1 (LMNB1) green fluorescent protein (GFP) reporter iPSC line^45^ and confocal microscopic analysis at 3 h post-passage, we observed irregularly shaped nuclei in the presence of DMSO and Y-27632, whereas CEPT treatment maintained normal circularity of iPSC nuclei (Fig. 7a). Importantly, under DMSO or Y-27632 treatment, phosphorylation of histone variant H2AX on serine 139 (γH2AX), a marker for DNA double-strand breaks and genomic instability^58^, was detected by immunocytochemistry and Western blotting. In contrast, γH2AX was absent in CEPT-treated cultured (Fig. 7b,g and Supplementary Fig. 19a). This striking finding suggests that DNA damage occurs more frequently during routine passaging of hPSCs than previously anticipated and can be prevented by CEPT. Next, immunocytochemical analysis of actin and non-muscle myosin (MYH10) revealed dramatic differences in cytoarchitecture at 3 h post-plating (Fig. 7c). Signs of cellular stress were detected in the presence of DMSO such as blebbing due to constant cell contractions^15^. As expected, treatment with Y-27632 prevented blebbing but led to the formation of actin stress fibers at the colony edge as previously reported^31^. Remarkably, cells readily attached to vitronectin-coated plates and displayed normal morphologies in the presence of CEPT (Fig. 7c).

**Figure 7.**
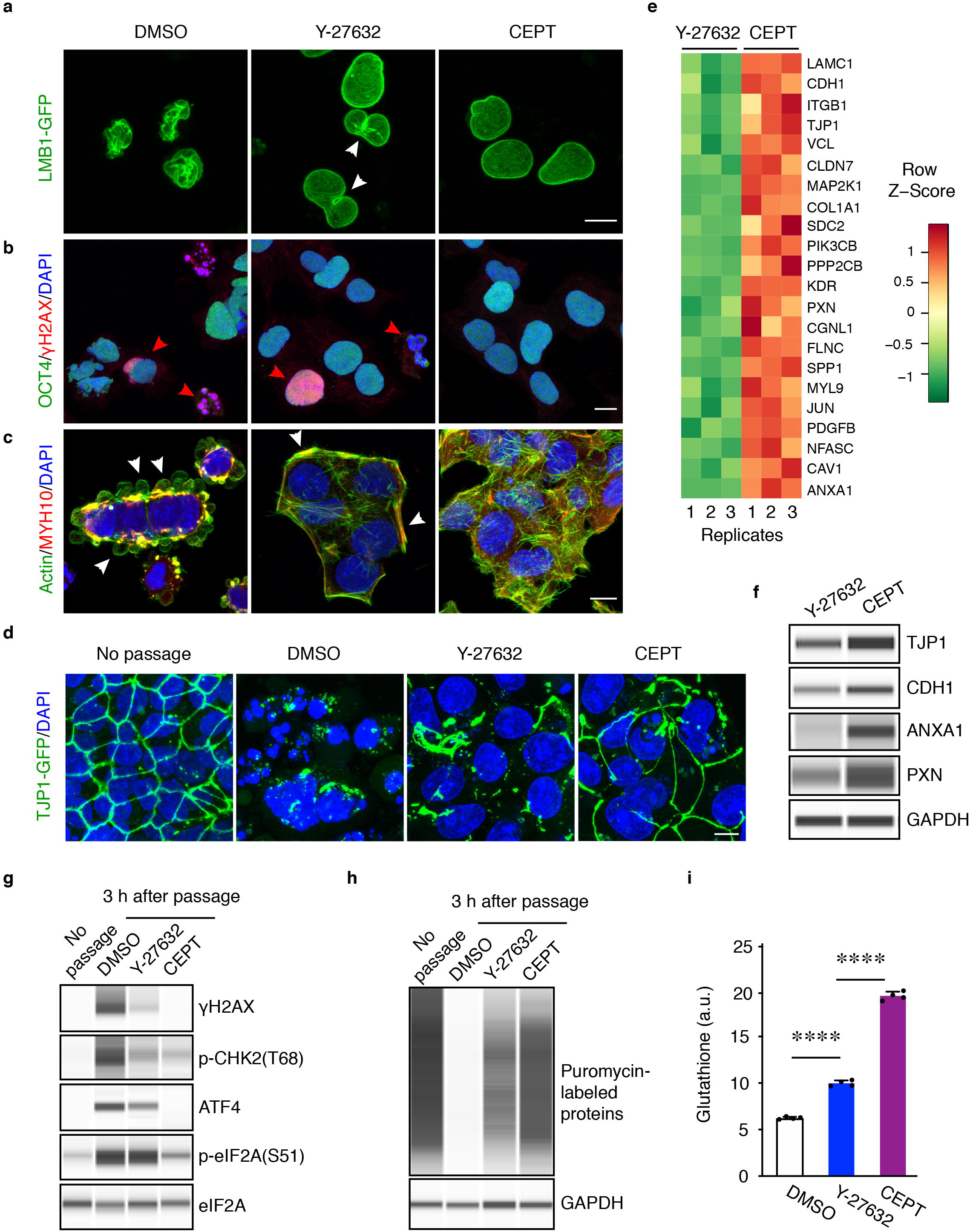
CEPT protects dissociated cells from multiple cellular stress mechanisms. **a,** Confocal microscopic analysis of the lamin B1-GFP reporter cell line displaying dramatic morphological differences in nuclear shape during cell passaging (30 min after plating). Note the abnormal nuclear morphology of cells undergoing contractions in the presence of DMSO. Although Y-27632 inhibits cell contractions and nuclear morphologies appear more normal, furrow-like constrictions (white arrowheads) indicate that nuclei are affected by ongoing physical stress. In contrast, cells exposed to CEPT maintain normal circular nuclei. Scale bar, 10 μm. **b,** OCT4 expressing cells were immunoreactive for γH2AX when exposed to DMSO and Y-27632 (red arrowheads) but not when treated with CEPT (3 h post-plating). Scale bar, 10 μm. **c,** Dramatic cytoskeletal differences during cell passaging (3 h post-plating) as measured by immunocytochemistry against actin and myosin. Stressed cells show blebbing (white arrowheads) in the presence of DMSO or form prominent actin stress fibers at the colony edge when exposed to Y-27632 (white arrowheads). Note the normal morphology and cell attachment in the presence of CEPT. Scale bar, 10 μm. **d,** Confocal analysis of an iPSC GFP reporter cell line visualizing the tight junction protein TJP1 (ZO-1) before passaging and after cell dissociation. Representative images were taken 3 h post-plating for DMSO, Y-27632, and CEPT. Note the dramatic differences across treatment groups and better recovery of dissociated cells when treated with CEPT. Scale bar, 10 μm. **e,** RNA-seq analysis of hESCs (WA09) treated with Y-27632 or CEPT for 24 h. **f,** Western blot analysis of hESCs (WA09) treated with Y-27632 or CEPT. Several membrane-associated proteins were expressed at higher levels after CEPT treatment (24 h post-plating). Similar results were obtained with iPSCs (see Supplementary Fig. 20). **g,** Western blot experiments of hESCs (WA09) showing strong stress response in cells treated with DMSO or Y-27632 (3 h post-passage). Note the absence of γH2AX and ATF4 in CEPT-treated cells resembling the control group (no passage). Similar results were obtained with iPSCs (see Supplementary Fig. 19a). **h,** Puromycin pulse-chase experiment of hESCs (WA09) demonstrate that protein synthesis was strongly impaired during cell passaging and can be rescued by CEPT (3 h post-passage). See also Supplementary Fig. 19b confirming all observations with iPSCs. **i,** Glutathione levels were significantly higher in hESCs (WA09) passaged with CEPT (3 h post-plating, n = 4 wells for each group). \*\*\**P* < 0.0001, one-way ANOVA. Similar results were obtained with several hESC and iPSC lines (see Supplementary Fig. 21).

To further characterize these marked differences, we performed live-cell imaging and monitored the expression of the tight junction protein TJP1 (also known as ZO-1) by using an iPSC GFP reporter line^45^. Again, significant morphological differences were evident between all groups at 3 h post-passage indicating that CEPT-treated cells were less stressed and recovered faster from single cell dissociation compared to DMSO and Y-27632 (Fig. 7d). Next, RNA-seq experiments were performed to analyze the transcriptome of cells treated with Y-27632 and CEPT for 24 h. We noticed a trend in that proteins that mediate cell-cell and cell-extracellular matrix (ECM) interaction were expressed at higher levels in the presence of CEPT (Fig. 7e). Western blots confirmed that TJP1 (ZO-1) and other cell membrane-associated proteins such as cell adhesion molecule CDH1 (also known as E-CADHERIN), focal adhesion molecule paxilin (PXN), and Annexin A1 (ANXA1; regulates rearrangement of actin cytoskeleton, cell polarization and migration) were expressed at higher levels in the presence of CEPT versus Y-27632 (Fig. 7f and Supplementary Fig. 20). Together, these observations suggest that CEPT protects cells from anoikis more efficiently than Y-27632.

To delineate other cytoprotective mechanisms provided by CEPT, we performed additional Western blot experiments at 3 h post-passage using hESCs and iPSCs. As mentioned above, DNA damage was indicated by γH2AX expression and could be prevented by CEPT (Fig. 7b,g). Phosphorylation of checkpoint kinase 2 (CHK2), a marker for cell cycle arrest^60^, was detectable at higher levels in cultures exposed to DMSO and Y-27632 versus CEPT (Fig. 7g and Supplementary Fig. 19a). These observations are important because DNA damage and re-entering the cell cycle after cell dissociation are bottlenecks for hPSC survival^9, 61^. Activating transcription factor 4 (ATF4) is a master regulator of cellular stress response in normal cells (e.g. unfolded protein response, ER stress, amino acid deprivation, hypoxia) and is frequently upregulated in cancer cells ^62^. Notably, CEPT-treated cells and controls (no passage) did not express ATF4 whereas cultures treated with DMSO or Y-27632 strongly induced ATF4 in hESCs as well as iPSCs (Fig. 7g and Supplementary Fig. 19a). Moreover, CEPT treatment resulted in significantly lower levels of phosphorylated serine 51 of eIF2A, whereas high phosphorylation levels in DMSO and Y-27632 indicated protein translation arrest^32^ (Fig. 7g and Supplementary Fig. 19a). Indeed, pulse-chase puromycin experiments ^63^ confirmed that protein synthesis was strongly impaired during cell passaging with DMSO, improved by Y-27632, but was markedly higher after CEPT treatment at 3 h post-passage (Fig. 7h and Supplementary Fig. 19b). Lastly, we measured glutathione levels after single cell dissociation. The tri-peptide glutathione is one of the most abundant intracellular antioxidants that protects mammalian cells from oxidative stress by maintaining redox homeostasis^64^. As confirmed in different hESC and iPSC lines, glutathione levels were significantly higher in cells passaged with CEPT compared to DMSO and Y-27632 (Fig. 7i and Supplementary Fig. 21), consistent with the notion that hPSCs were less stressed and better protected by CEPT.

In summary, these findings provide evidence that CEPT buffers cellular stress and provides comprehensive cytoprotection, which is not achieved by currently used strategies. The multiple advantages of using CEPT include the following mechanisms: inhibition of cell contraction, maintenance of normal nuclear circularity, protection from DNA damage, prevention of cell cycle arrest, restoration of protein translation, higher expression level of proteins that enable cell-cell and cell-ECM interactions, and higher glutathione levels (Fig. 8a). Therefore, CEPT safeguards cell survival and clonal growth by favorably regulating complex interconnected mechanisms of cell structure and function that are otherwise compromised due to cascading events of cellular stress. In conclusion, CEPT represents a versatile platform that can broadly leverage multiple areas in basic research and regenerative medicine (Fig. 8b).

**Figure 8.**
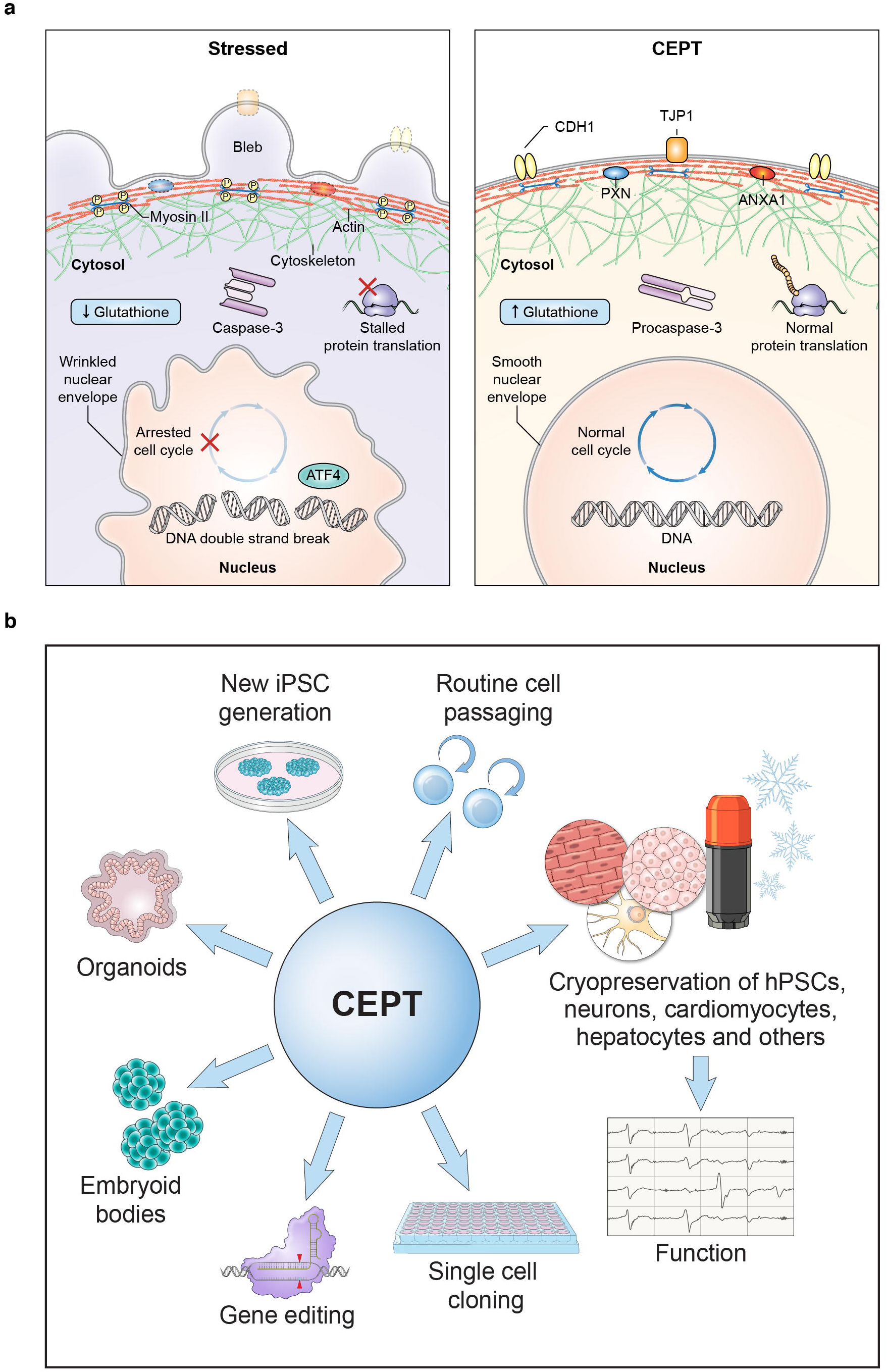
Comprehensive cytoprotection and versatile applicability of CEPT. **a,** Summary of the various interconnected stress mechanisms investigated in this study. Comparison between stressed (DMSO or Y-27632) and cytoprotected (CEPT) cells revealed dramatic differences affecting cell structure and function. Mitigation of complex stress mechanisms and conferring cell fitness by small molecule combinations such as CEPT may become a common strategy in cell biology and regenerative medicine. **b,** Overview of the various applications that were optimized by CEPT in the present study.

## Discussion

Controlling viability, safety, and cell function are key challenges for translational stem cell research. We addressed these challenges by performing iterative single-agent and combination screens in high-throughput 1536-well format using hPSCs (e.g. ultra-low cell density screening, full dose-response curves for nearly 16,000 compounds, identification of synergistic activities). We identified a small molecule cocktail that dramatically improves the culture and utilization of pluripotent as well as differentiated cells. Mechanistic follow-up studies revealed the importance of using a multi-target polypharmacology strategy for cytoprotection and maintenance of normal cell structure and function during critical periods of cellular stress (Fig. 8a). The demonstration that CEPT was beneficial in widely used stem cell procedures (cell passaging, cryopreservation of undifferentiated and differentiated cells, improved cell function after thawing, EB and organoid formation, single cell cloning, genome editing, iPSC line establishment) has immediate and far-reaching implications for basic research and clinical applications (Fig. 8b). We propose that the versatility of CEPT will help to establish new quality control standards, increase experimental reproducibility, and bring the iPSC technology to patients more efficiently. Moreover, we anticipate that CEPT will become a more broadly used strategy in various other fields that currently rely on ROCK pathway inhibition such as reproductive biology, cryobiology, transplantation medicine, and cancer research (e.g. establishing new cancer cell lines from patient material, isolating cancer stem cells, producing chimeric antigen receptor T cells)^65, 66^.

Non-random genetic variations and oncogenic mutations can arise in hPSC cultures^67–69^. These changes occur largely for unknown reasons but increased passage number and cell dissociation correlate with the risk of accumulating genetic aberrations^9, 70^. Our data provide a new perspective by demonstrating that under currently used culture conditions, even during routine cell passaging, self-renewing hPSCs are exposed to unexpectedly high levels of stress affecting various cellular compartments. We propose a model whereby repeated exposure to stress leads to genetic abnormalities compromising normal fitness of self-renewing pluripotent cells with increased passage number. In contrast, CEPT promotes cytoprotective cell survival by safeguarding hPSCs against several stress mechanisms that require an integrated strategy for optimal outcome. The strong and highly reproducible optimizations, quantitatively and qualitatively, achieved here by CEPT provide a new rationale for culturing hPSCs in a safer and more efficient fashion and will help to develop advanced strategies for drug discovery, disease modeling, and personalized cellular therapies.

## Supplementary Figure Legends

**Supplementary Figure 1.**
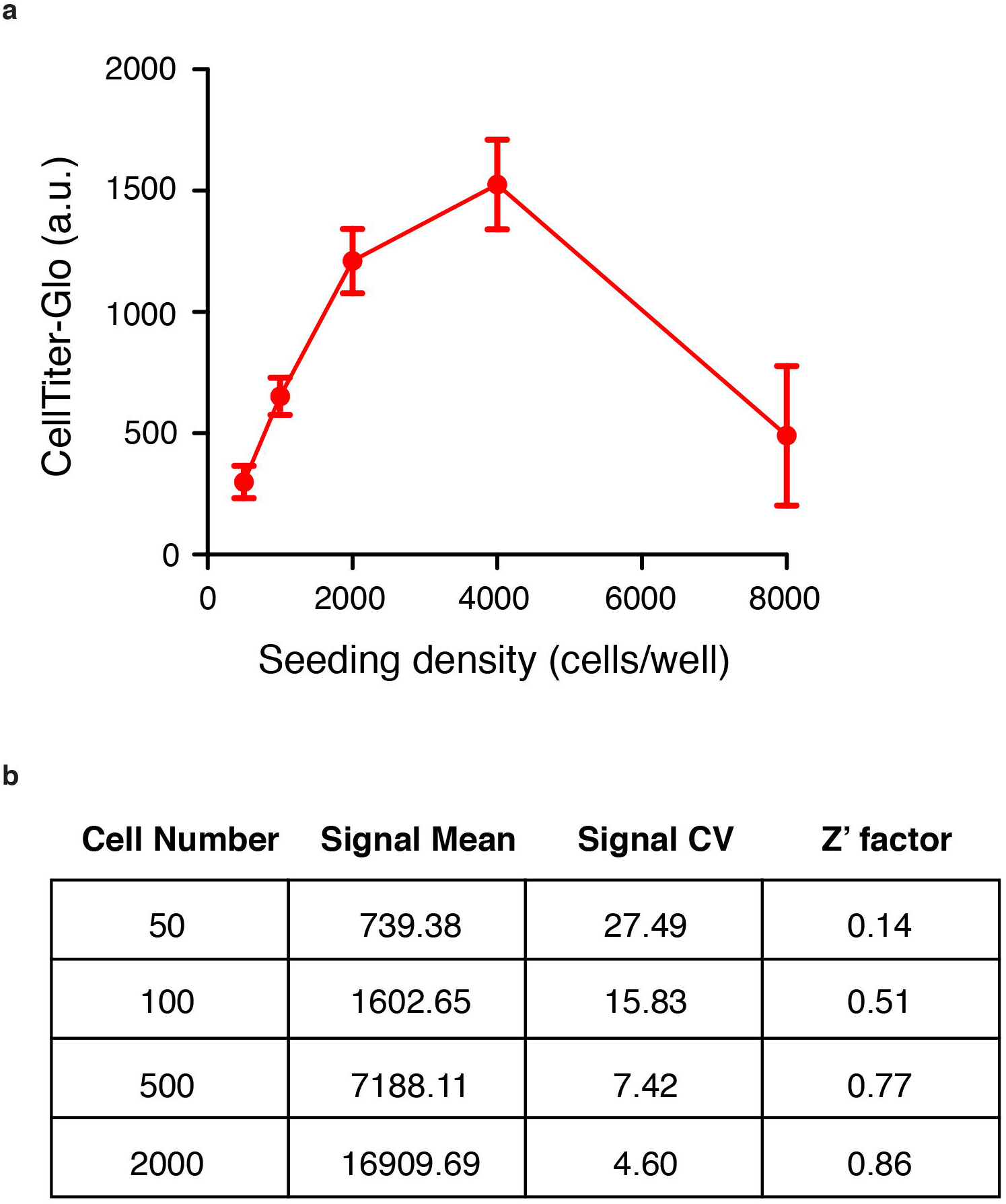
Assay development for cell viability screening in 1536-well format. **a,** Different numbers of hESCs (WA09) were dispensed into each well of 1536-well plates in the presence of 10 μM Y-27632. CellTiter-Glo (CTG) reagent was dispensed 24 h later to assess the number of live cells in each well (n = 128 wells for each group). **b,** Plating 500 cells/well was determined to provide an acceptable Z’ factor suitable for qHTS (n = 256 wells for each group).

**Supplementary Figure 2.**
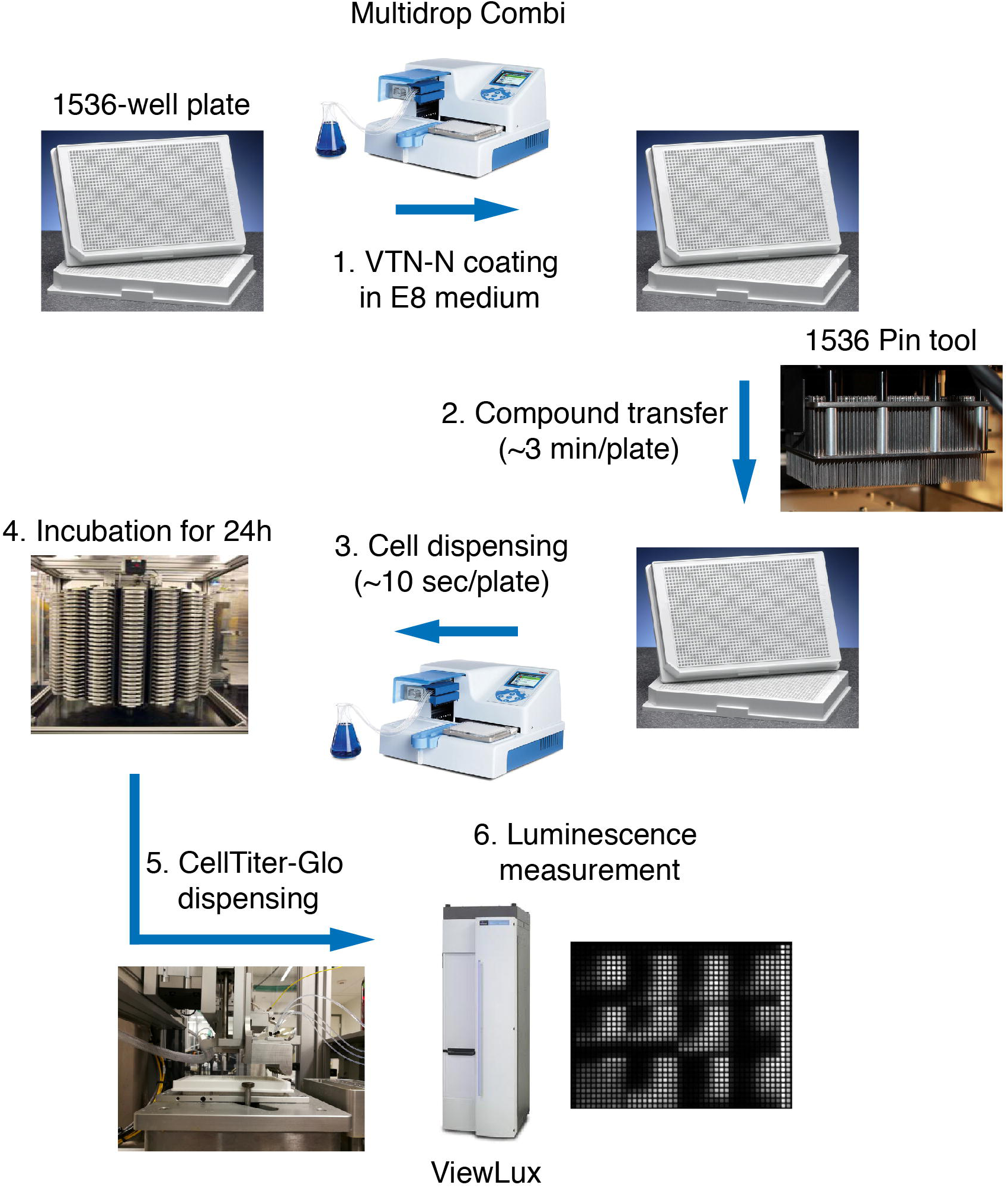
Workflow for qHTS in 1536-well plates optimized for hPSCs. Plates (1536-well format) were coated with VTN-N diluted in E8 medium at 37 °C for 1 h, followed by transfer of small molecule compounds (diluted in DMSO) from the compound library source plates by using a pintool. Once 1536-plates were ready, hESCs were dissociated with Accutase, re-suspended in E8 medium, and dispensed into the plates at a density of 500 cells/well. After the plates were incubated for 24 h, CTG reagent was dispensed and the luminescence signal was determined using the ViewLux plate reader.

**Supplementary Figure 3.**
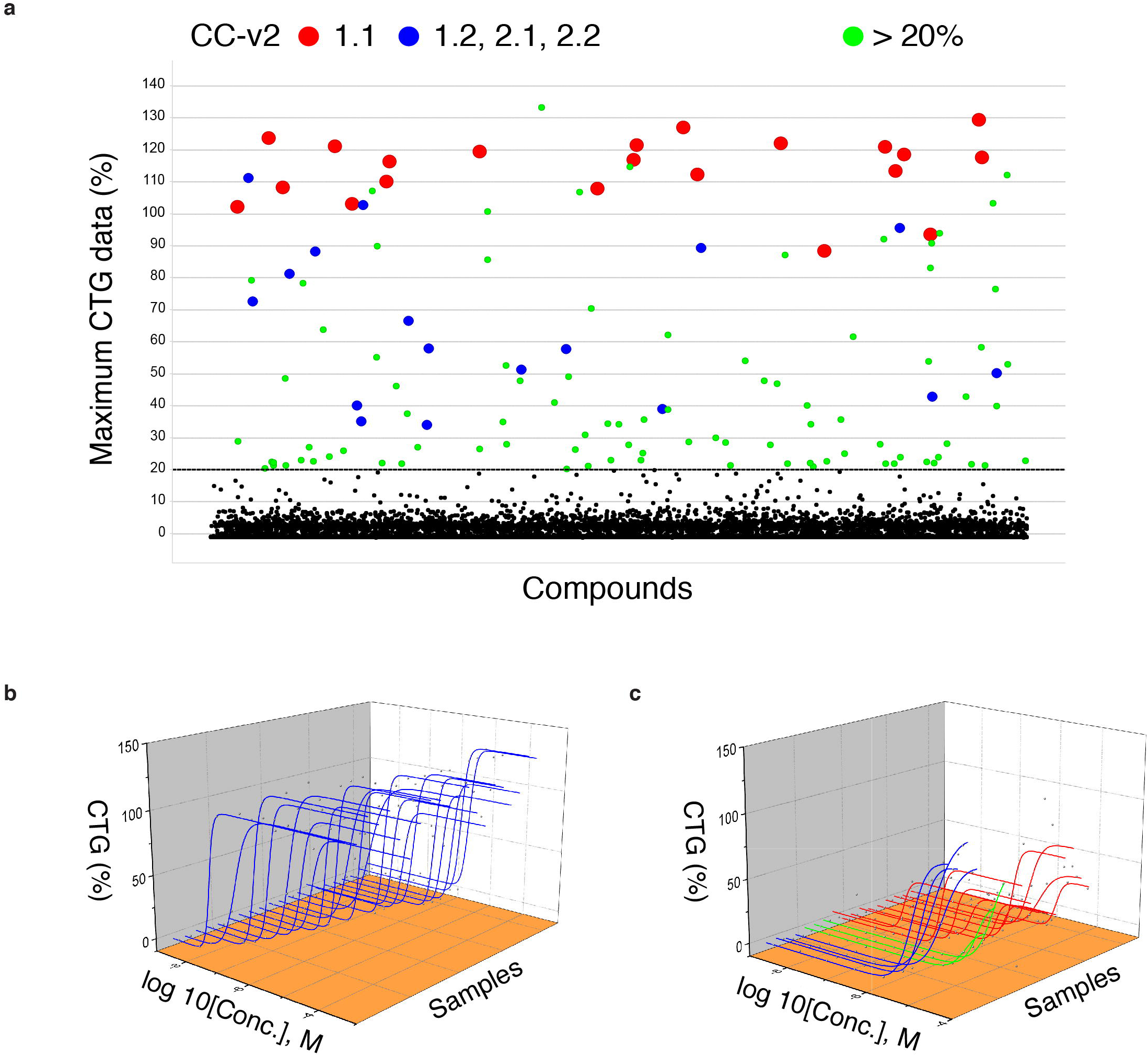
Overview and dose-response analysis of qHTS data. **a,** Scatter plot of maximum CTG values obtained for all compounds screened. Data were normalized to the CTG values obtained using 10 μM Y-27632, which was set as 100% (control). Setting a threshold of 20% resulted in 113 active compounds (red, blue and green circles). b-c, Compound dose-response curves classified as class 1.1 (b and indicated as red circles in panel a), and classes 1.2, 2.1, and 2.2 (c and indicated as blue circles in panel a) according to Huang, R. *et al.* ^28^.

**Supplementary Figure 4.**
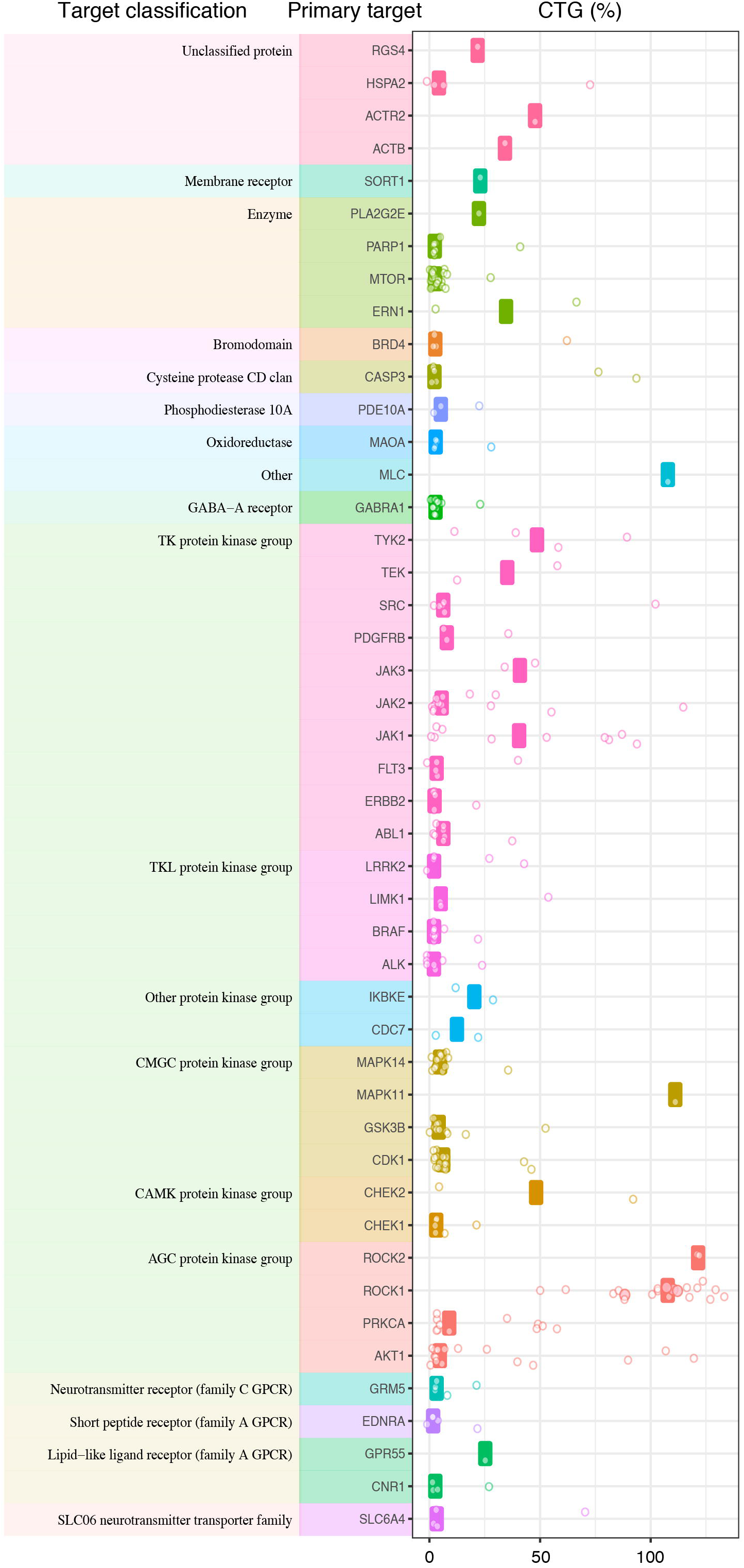
Target-based analysis of 113 hits discovered in the primary screen. Maximum cell survival achieved by the compounds in the primary screening binned per mechanistic classes. Maximum survival was normalized to the CTG reading obtained with 10 μM Y-27632 treatment. Each point represents a compound which has the designated primary target and target class on the left panel. The bars in the scatter plot are the median of maximum survival for the given target.

**Supplementary Figure 5.**
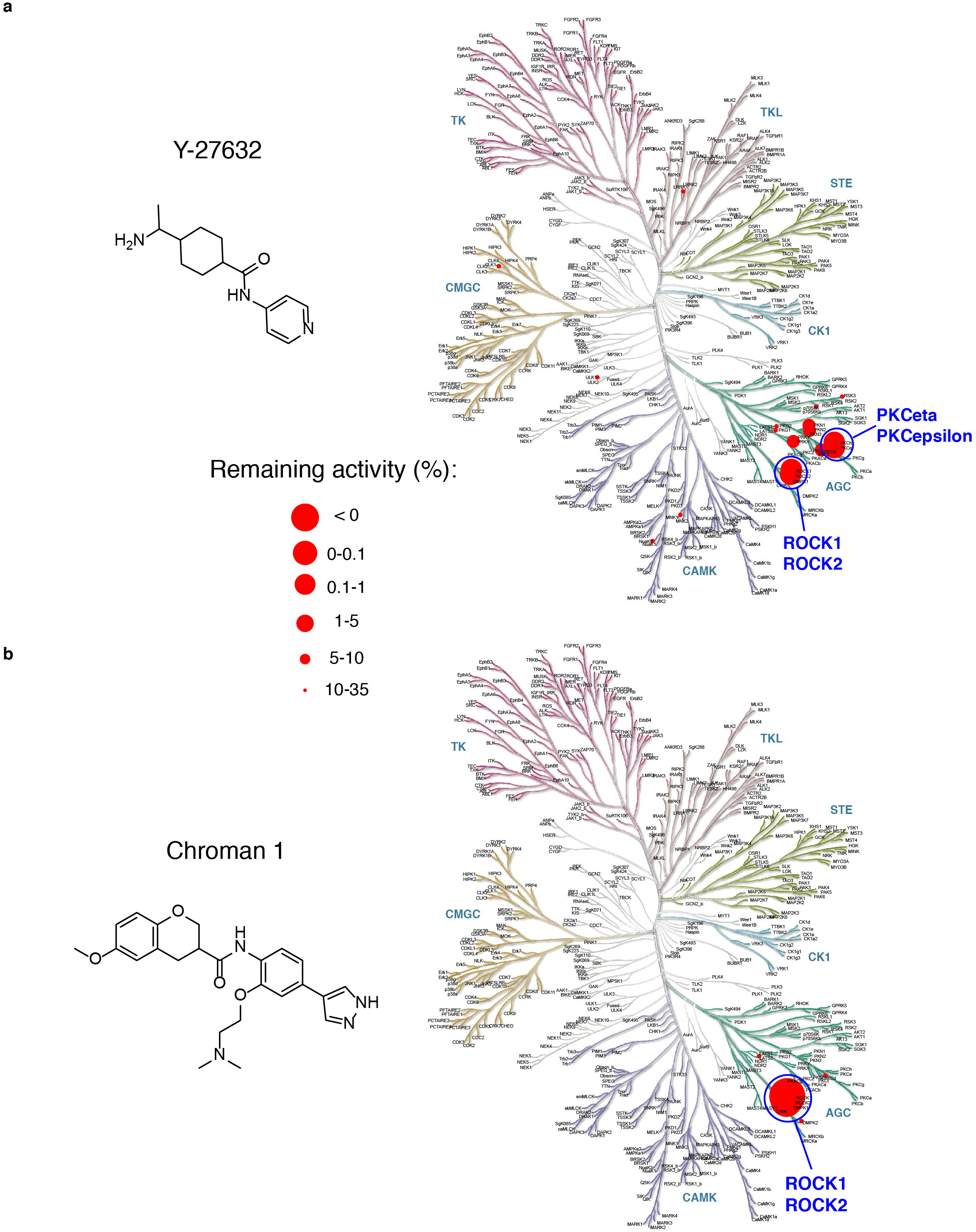
Superior target specificity of Chroman 1 as compared to Y-27632. HotSpot kinase profiling was used to individually inhibit a panel of 369 human wild-type kinases using 50 nM Chroman 1 and 10 μM Y-27632 (concentrations were chosen based on dose-response curves and maximum cell survival in the primary screen). Note the significant differences among these two ROCK inhibitors revealing that Chroman 1 is more potent and has fewer off-targets than Y-27632. See also Supplementary Fig. 6 for more details.

**Supplementary Figure 6.**
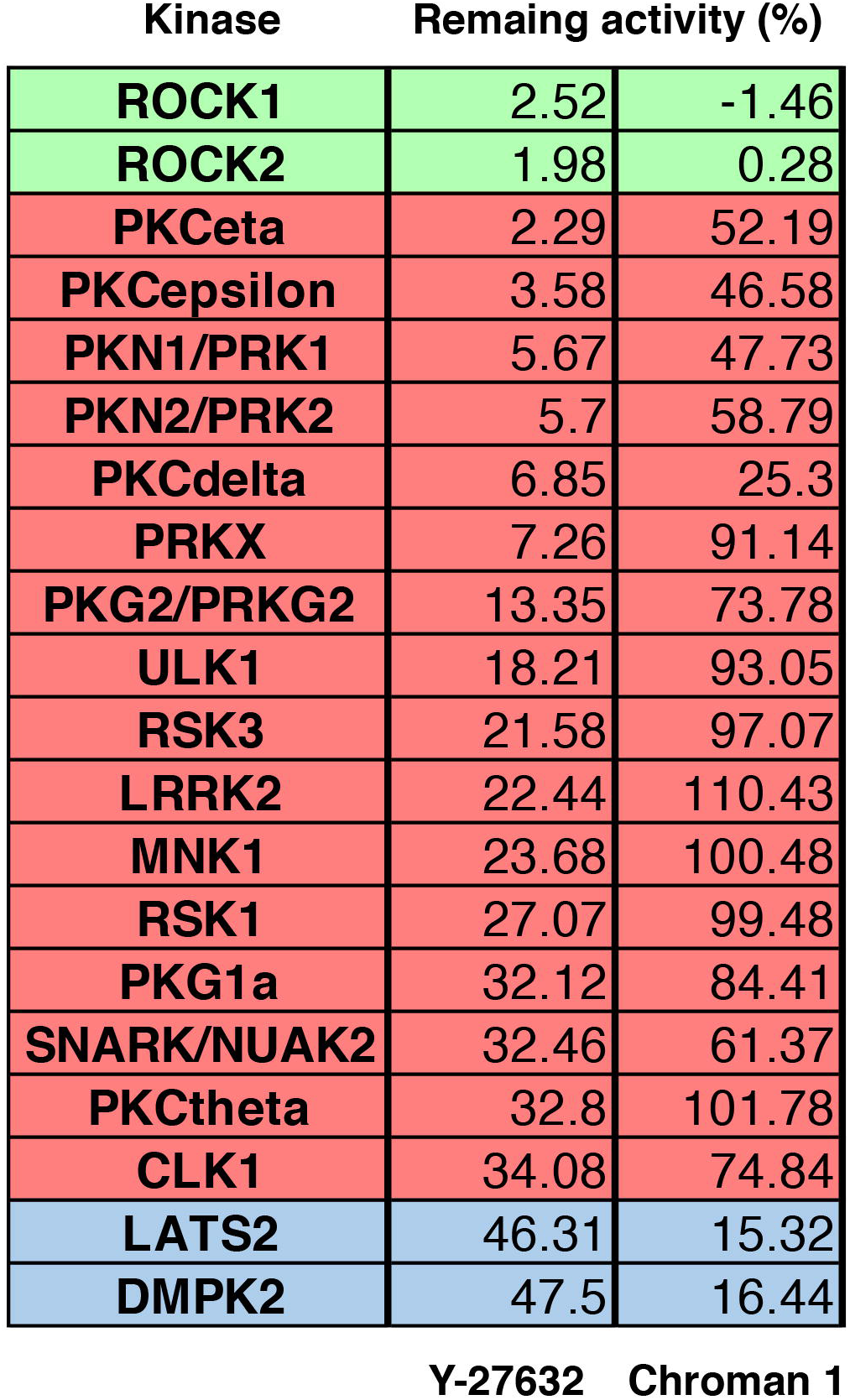
Inhibitory activity of Chroman 1 and Y-27632 measured by HotSpot kinase profiling. Table summarizing the inhibitory activity of 50 nM Chroman 1 and 10 μM Y-27632. Green portion of the table shows values for ROCK1 and ROCK2 as the primary targets, red portion indicates all off-target kinases that Y-27632 inhibits more strongly than Chroman 1. These include PKCeta (also known as PKCη or PRKCH), PKCepsilon (also known as PKCε or PRKCE), PKCdelta (also known as PKCδ or PRKCD), PKN1, PKN2 and PRKX. The only two off-target kinases that Chroman 1 inhibits relatively better than Y-27632 are LATS2 and DMPK2 (blue portion). Importantly, although numbers indicate a trend in this comparative assay, only values under 10% represent significant inhibitory activity^30^. See also Supplementary Fig. 5.

**Supplementary Figure 7.**
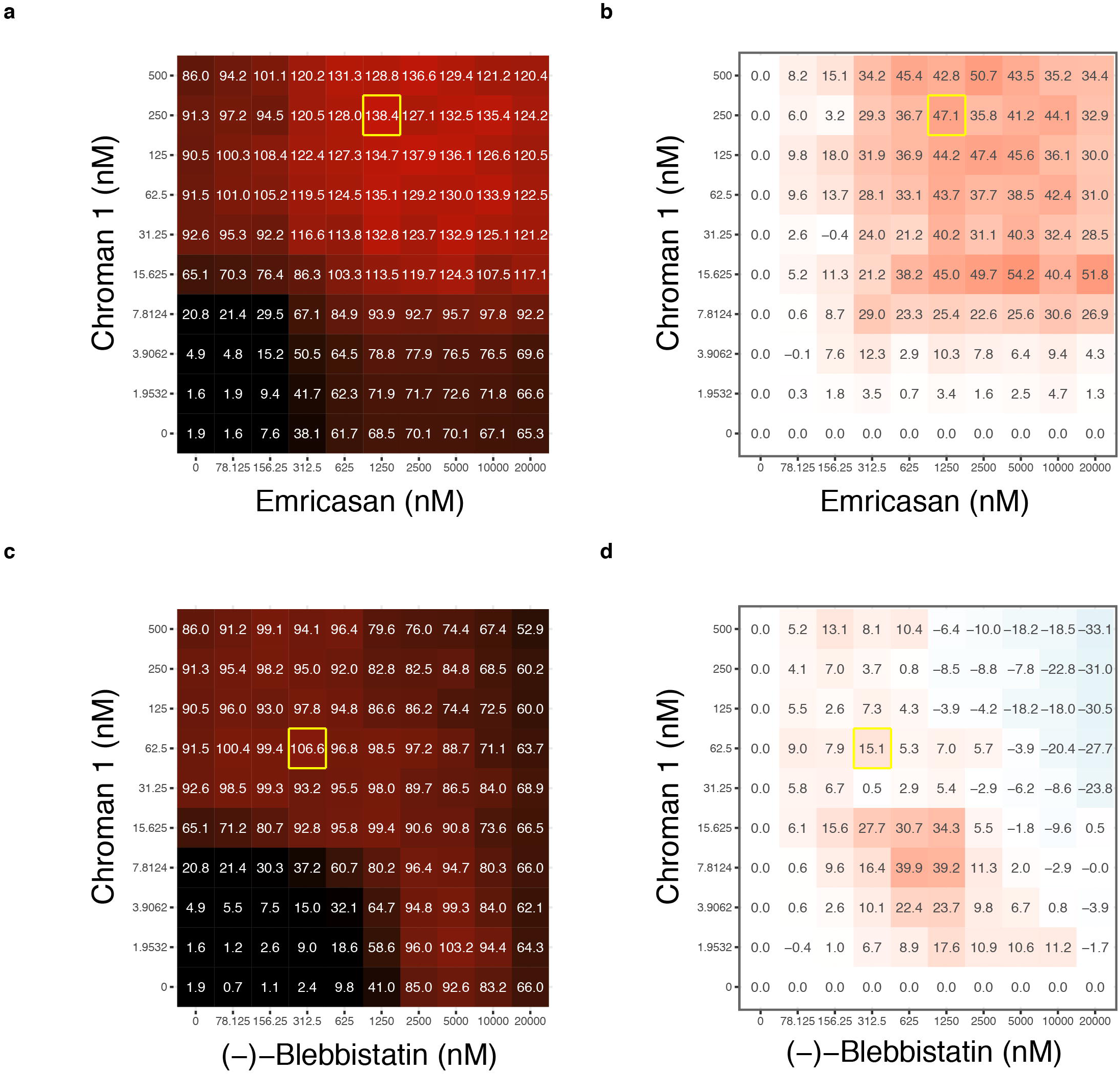
Checkerboard examples from the matrix screening data. The 10 x 10 viability matrices (**a** and **c**) and the delta HSA value matrices (**b** and **d**) for the combination of Chroman 1 + Emricasan (**a** and **b**), and Chroman 1 + Blebbistatin (**c** and **d**). Chroman 1 + Emricasan enhanced the maximum survival compared to Chroman 1 or Emricasan alone, which indicates synergistic activity. In contrast, Chroman 1 + Blebbistatin failed to further improve cell survival. Yellow boxes highlight the maximum survival and synergy level achieved by the combinations.

**Supplementary Figure 8.**
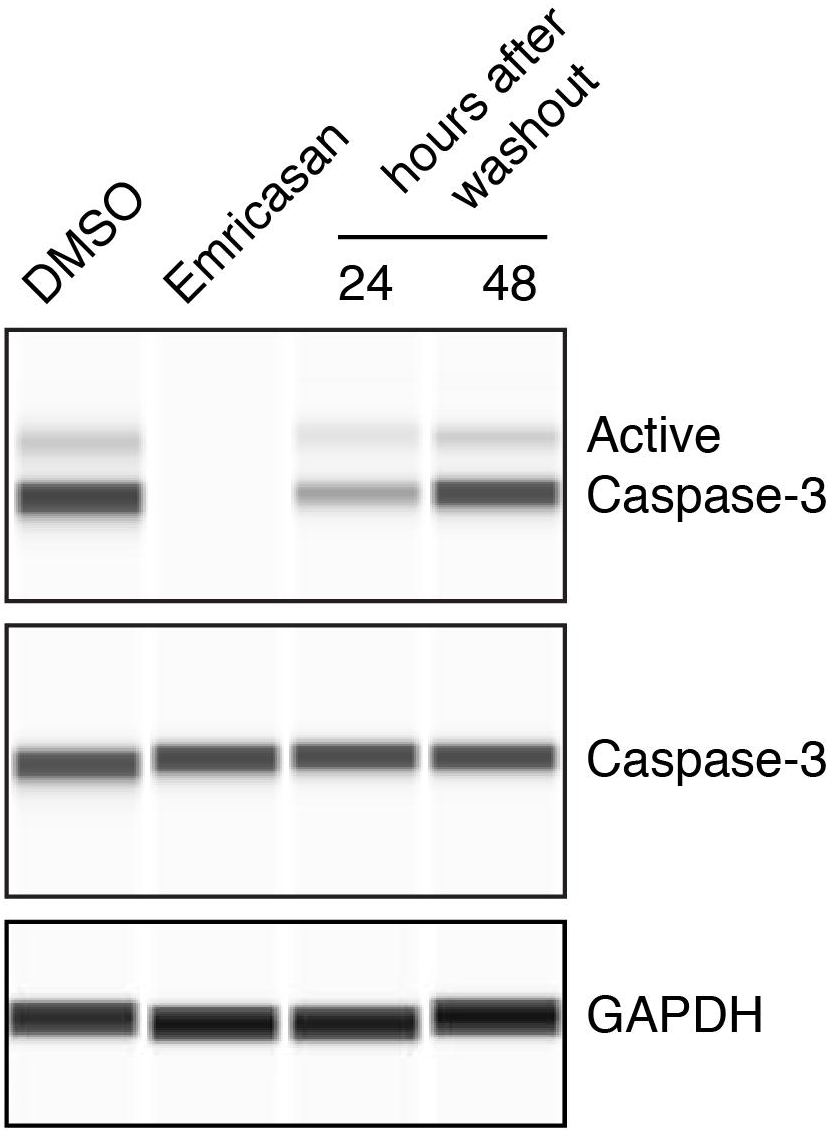
Reversible effects of Emricasan on Caspase-3 activation after cell dissociation. Emricasan efficiently blocked the cleavage and activation of Caspase-3 induced by hESC dissociation but did not impact total Caspase-3 levels. Note that active Caspase-3 returned to basal expression levels 24-48 h after Emricasan removal.

**Supplementary Figure 9.**
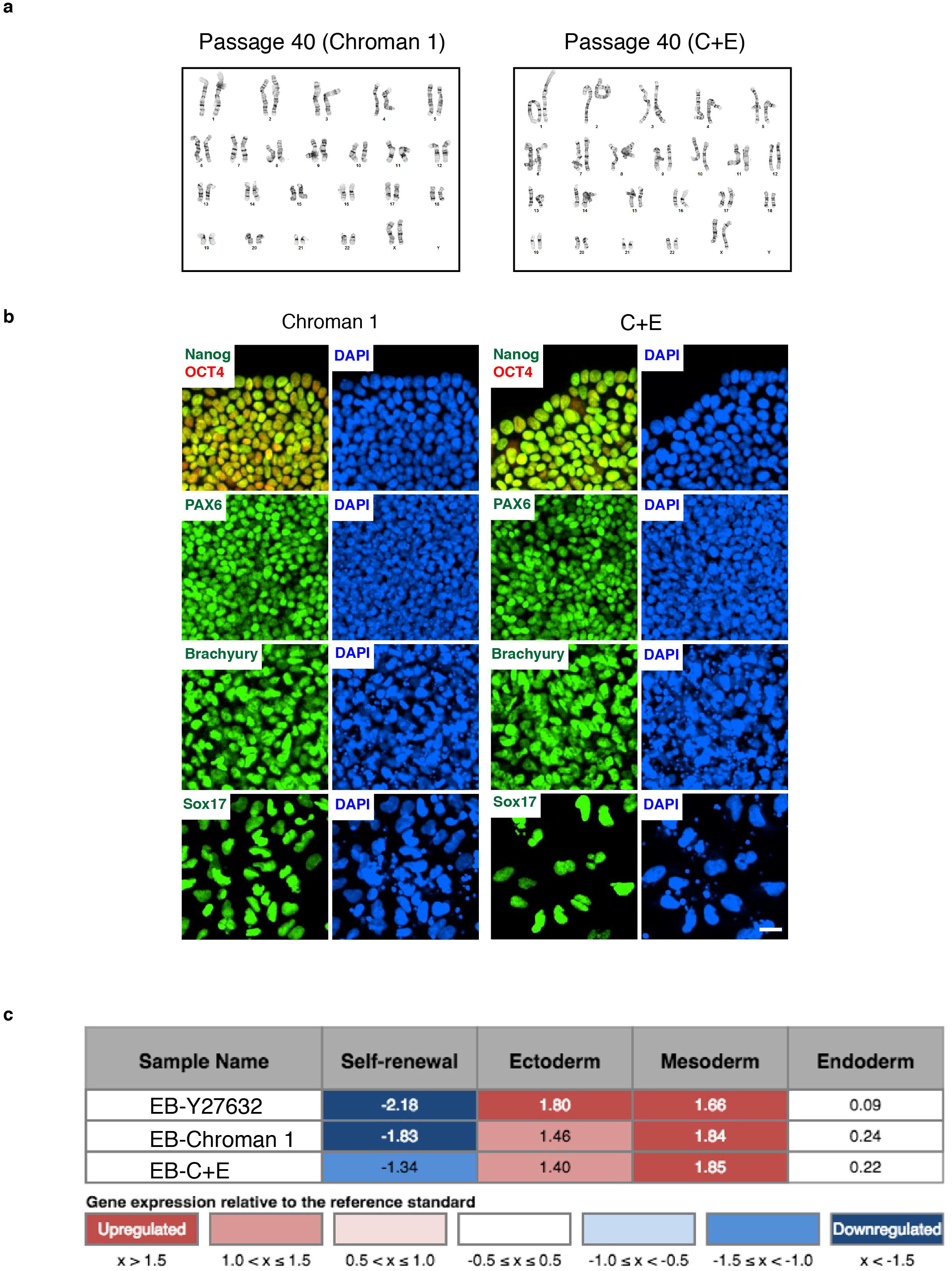
Repeated 24 h treatment with Chroman 1 or C+E for 40 passages. hESCs (WA09) were cultured over 40 passages with Chroman 1 or C+E applied for the initial 24 h during every passage. Cells maintained normal karyotypes (**a**), expressed pluripotent genes including NANOG and OCT4, and differentiated into ectoderm, mesoderm and endoderm lineages both by directed differentiation in adherent cultures (**b**) and spontaneous differentiation in EB cultures followed by ScoreCard analysis (**c**). Scale bar in **b**, 2.5 μm.

**Supplementary Figure 10.**
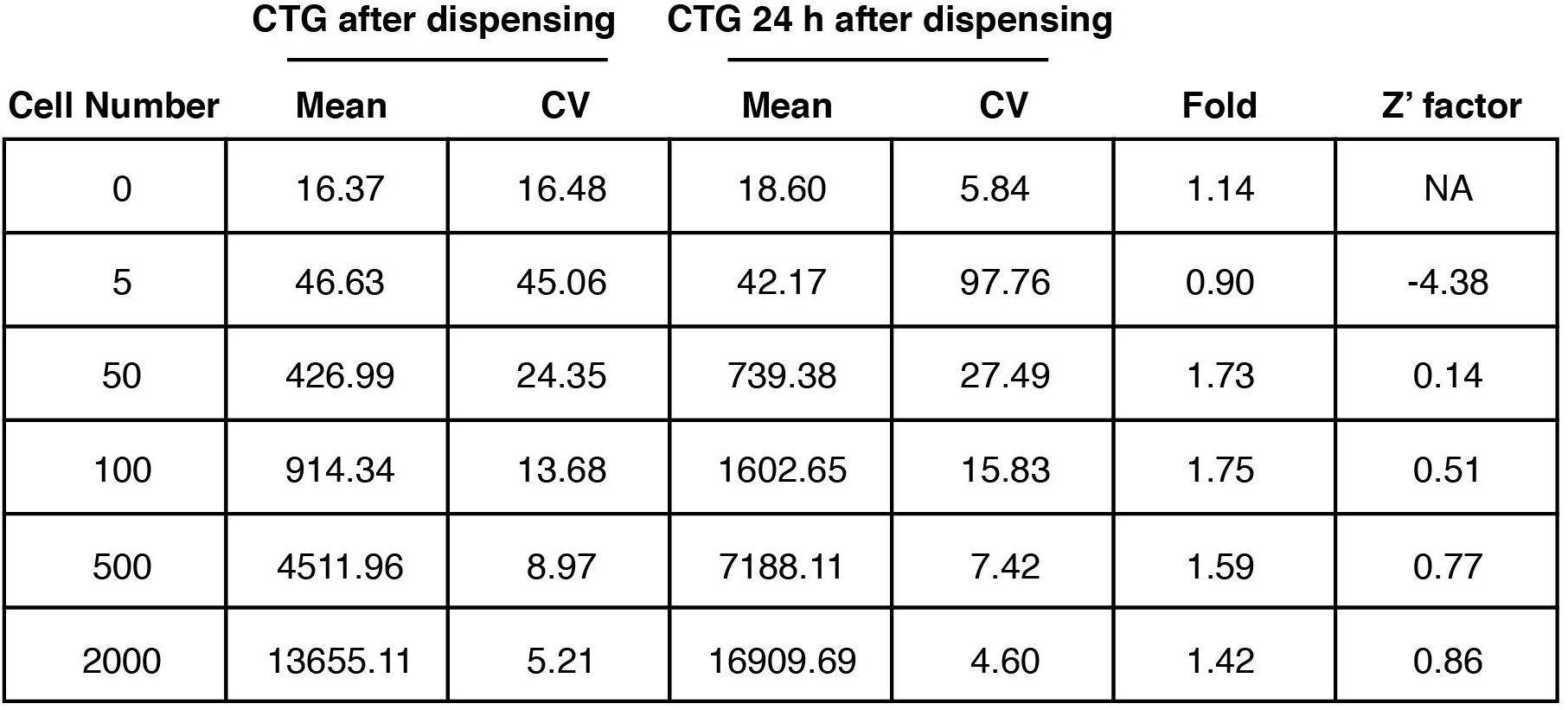
Assay development to determine hPSC survival and growth at different cell seeding densities in 1536-well format. hESCs (WA09) were single-cell dissociated with Accutase and dispensed into 1536-well plates at varying densities. Two plates were prepared for each seeding density. CTG reading was carried out with one plate immediately following cell dispensing (read 1) and with the second plate 24 h later (read 2). CTG-fold change was calculated as read 2 divided by read 1 to indicate cell recovery and growth within 24 h. Note the biphasic relationship between cell seeding density and the CTG-fold change, indicating that both high cell density (2000 cells/well) and ultra-low cell density (< 50 cells/well) impeded cell survival. (n = 128 wells for 0 and 5 cells/well, and n = 256 wells for 50, 100, 500 and 2000 cells/well).

**Supplementary Figure 11.**
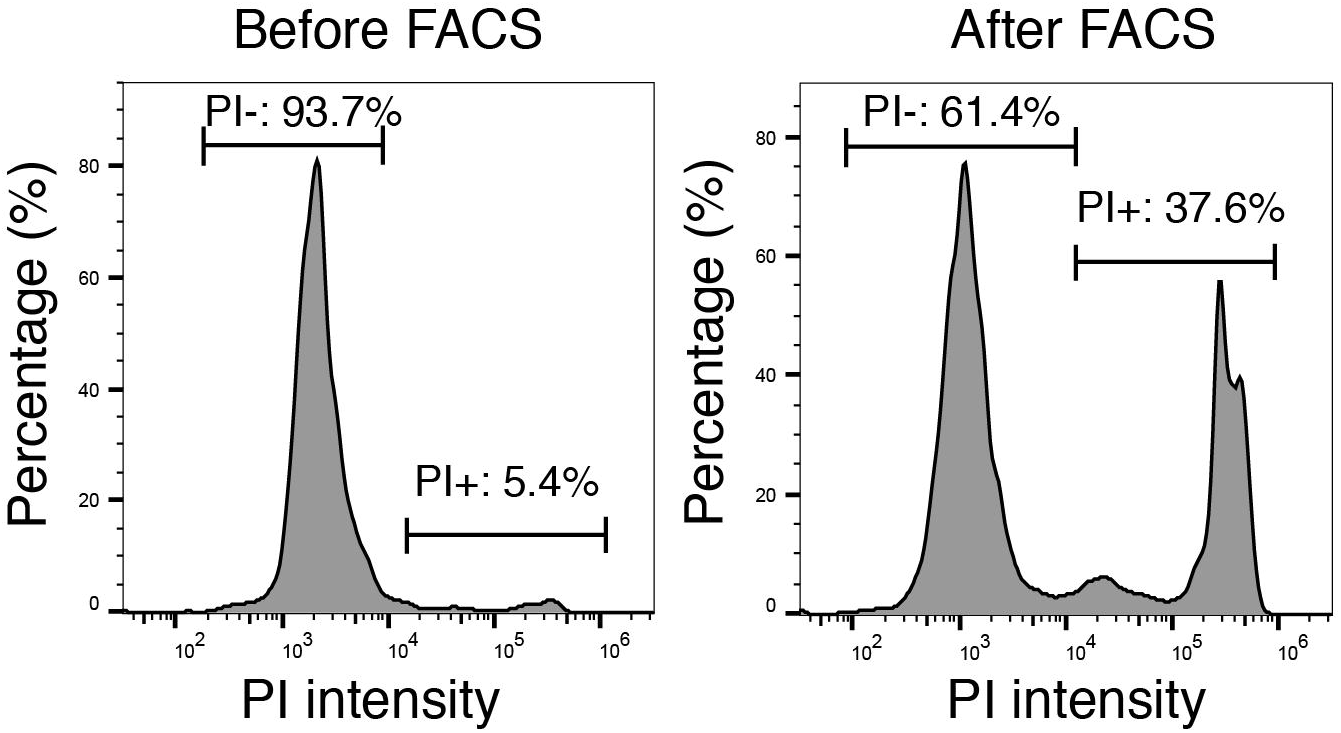
Vulnerability of hPSCs to the FACS sorting process. hESCs (WA09) were dissociated into single cells using TrypLE, and cell viability was assessed using PI immediately following cell dissociation and after FACS sorting.

**Supplementary Figure 12.**
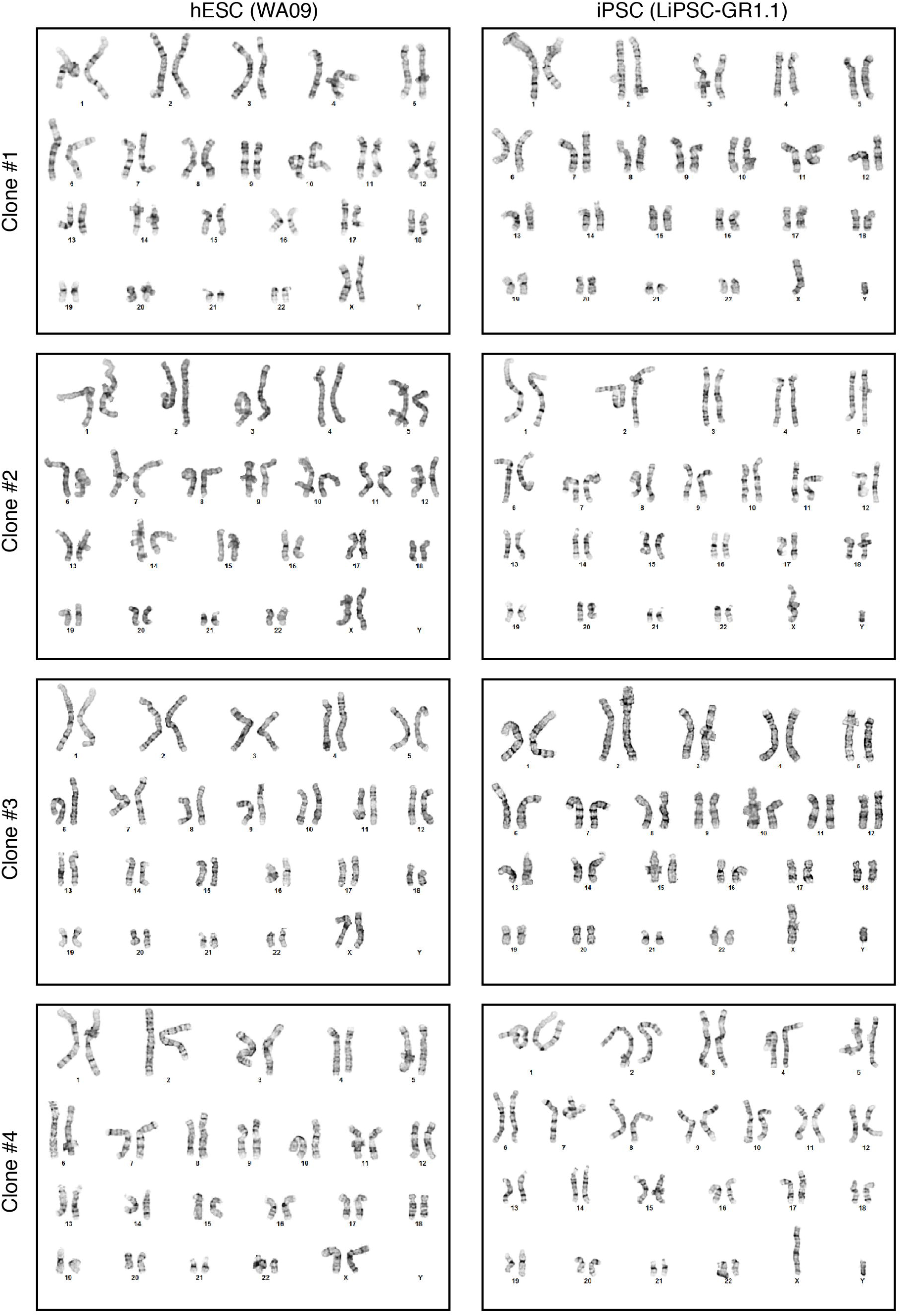
Normal karyotype of single-cell clones derived with CEPT treatment. After cell dissociation, single cells were FACS-sorted into 96-well plates and treated with CEPT. Single clones were then picked and again expanded using CEPT for 24 h at each passage. This strategy allowed to rapidly establish 4 clonal cell line from hESCs (WA09) and 4 clonal cell lines from iPSCs (LiPSC-GR1.1). All clonal cell lines were karyotypically normal.

**Supplementary Figure 13.**
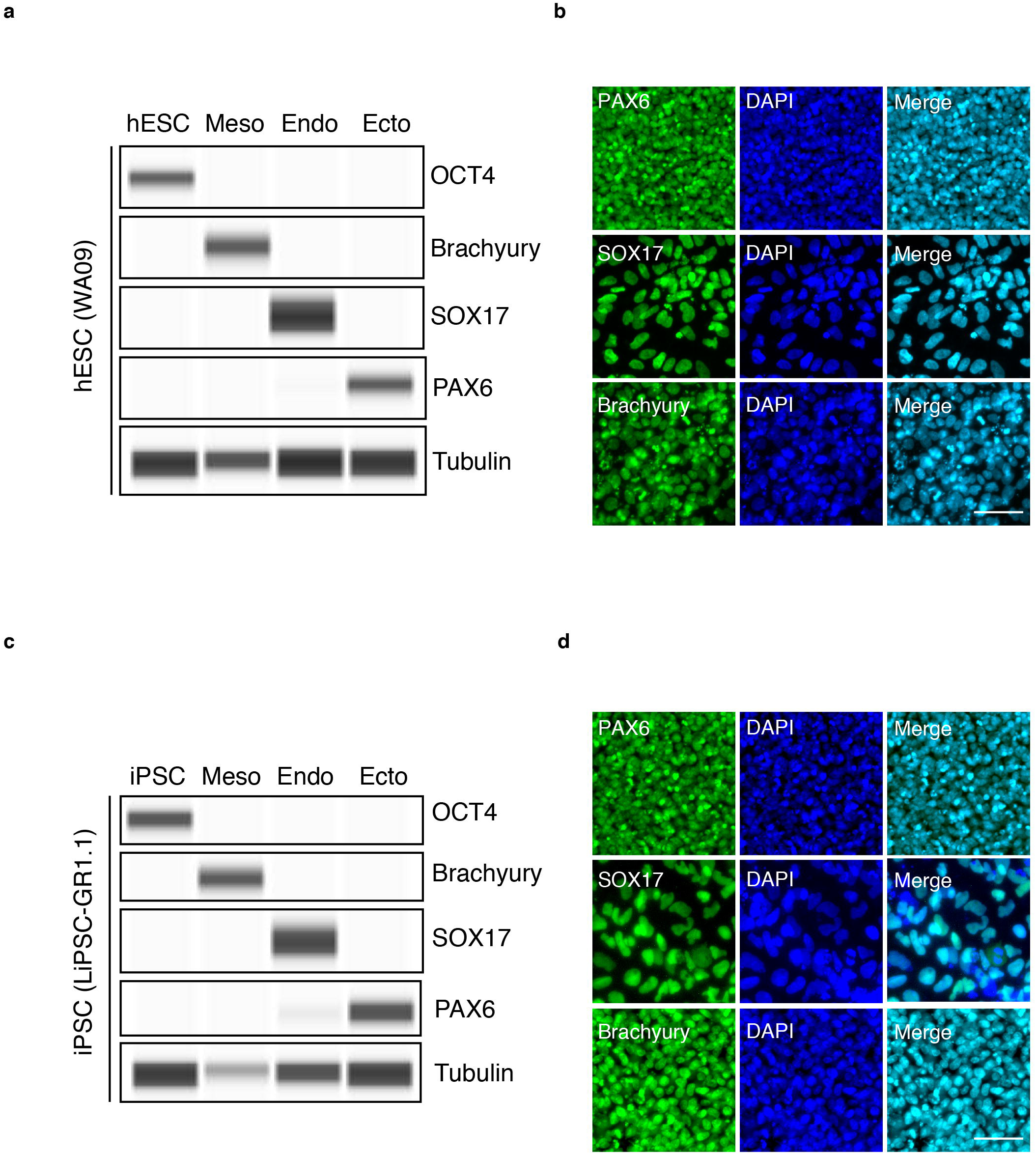
Maintenance of normal differentiation potential of hESCs and iPSCs after CEPT treatment. **a-c,** To demonstrate that CEPT treatment was safe, hESCs (WA09) and iPSCs (LiPSC-GR1.1) were cultured for 20 passages and exposed to CEPT for 24 h during every passage. Both cell lines maintained normal karyotypes, expressed OCT4 (Western blots in **a** and **c**), and differentiated into ectoderm, mesoderm and endoderm lineages (Western blot and immunocytochemistry **a** to **d**) by directed differentiation in adherent monolayer cultures. Scale bars: 50 μm.

**Supplementary Figure 14.**
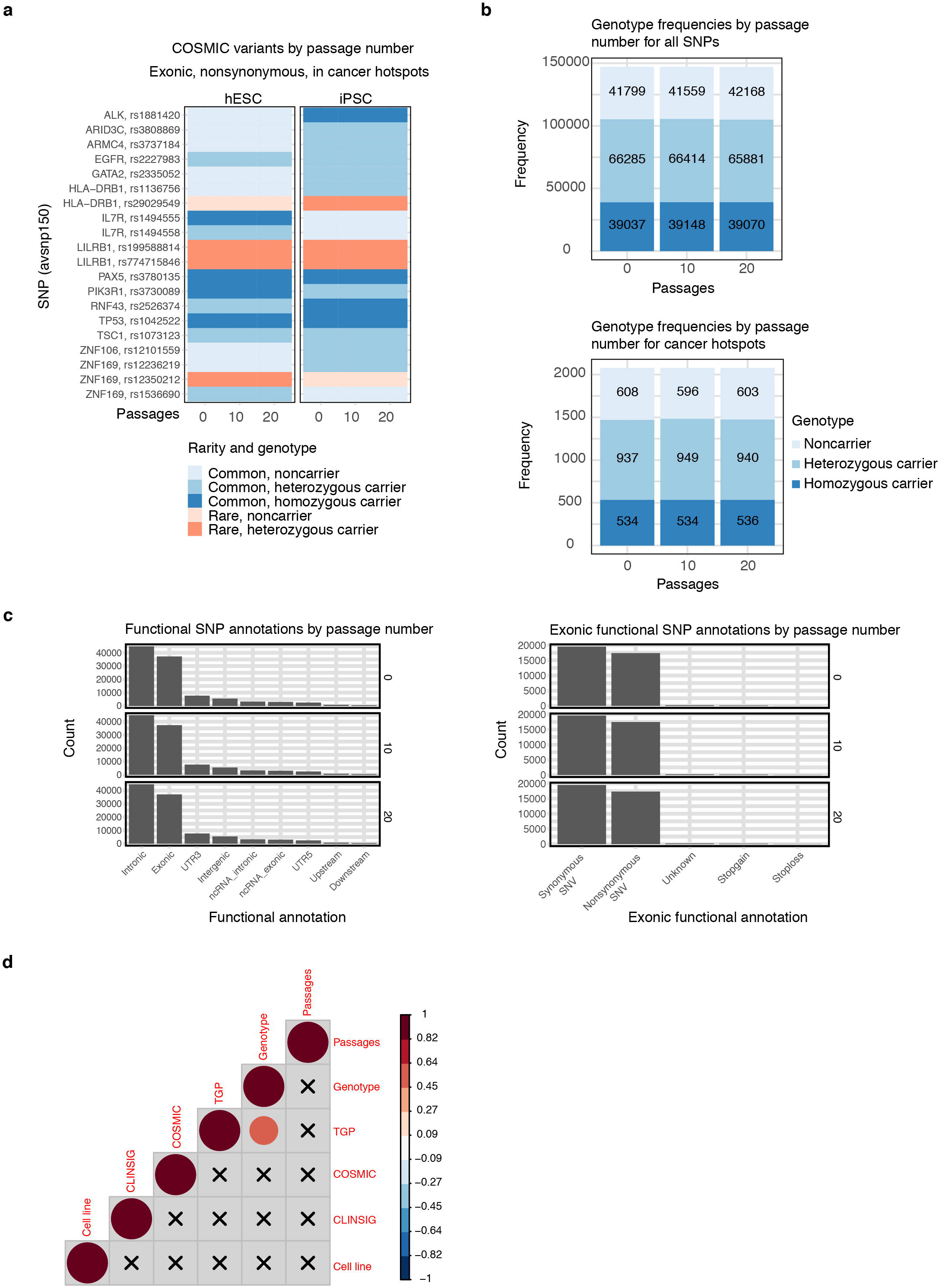
WES variant annotations, genotypes, and CNV analysis show genetic stability of cell lines. **a**, COSMIC variants by passage number for only exonic, nonsynonymous SNPs in cancer hotspots, split by cell line. There was no change in rarity or genotype over passaging. **b,** Genotype frequencies by passage number for all SNPs (upper panel) and SNPs in cancer hotspots (lower panel), which were proportionally constant. **c,** Functional SNP annotations by passage number (left) and exonic functional SNP annotations (right), a subset of the former, with unchanging variant counts per annotation category. **d,** Indel correlogram showing no significant correlation between key variables, passage number, and genotype. The CLINSIG database was used to analyze indels.

**Supplementary Figure 15.**
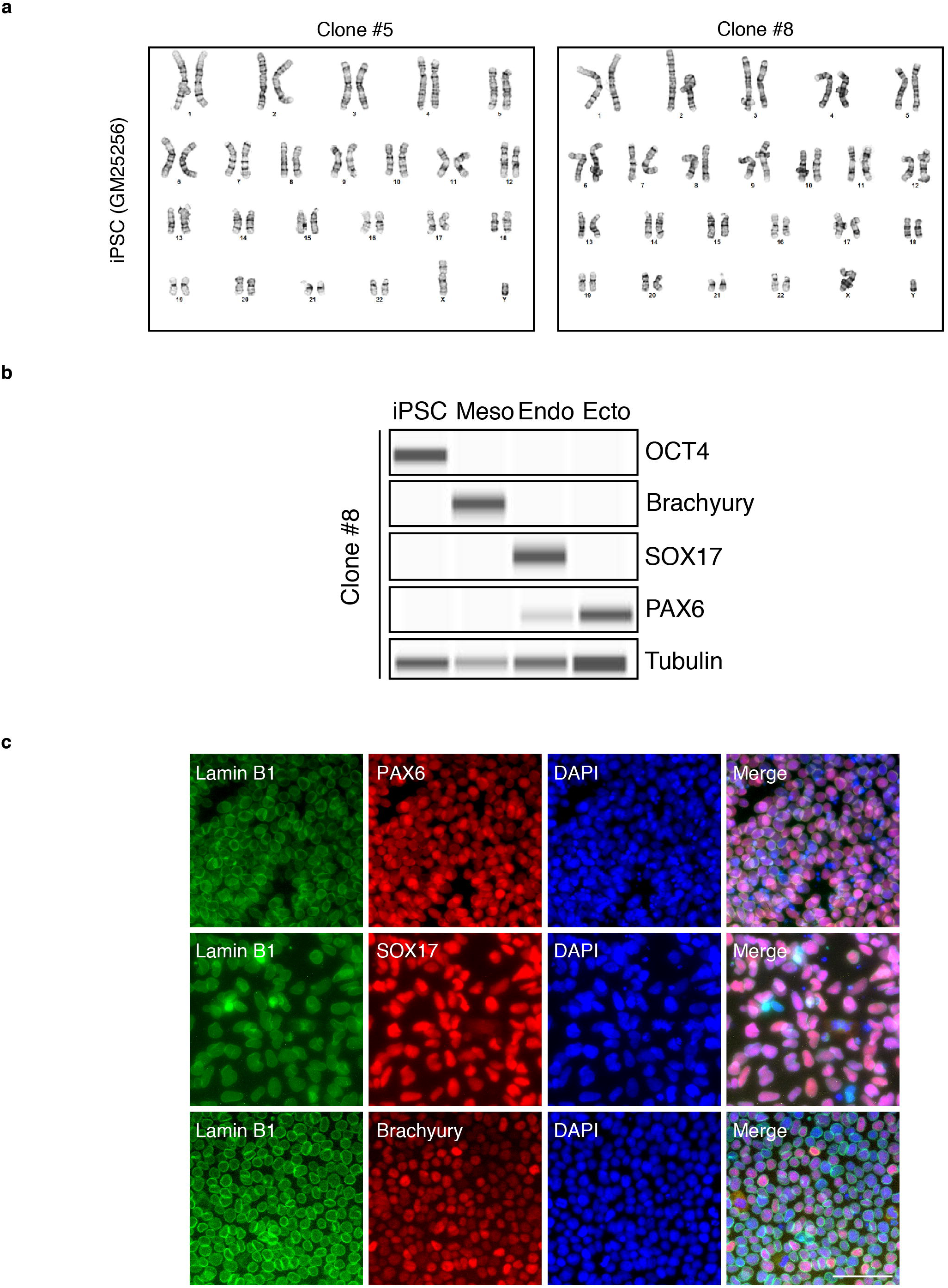
Normal karyotype and differentiation potential of LMNB1-edited iPSC clones. **a,** After gene editing, LMNB1-edited clones #5 and #8 were established as cell lines, expanded, and their normal karyotype confirmed. **b-c,** Western blot and immunocytochemical analysis of gene-edited and clone cell line demonstrating expression of typical markers for pluripotent (OCT4) and differentiated cells (Brachyury for mesoderm; SOX17 for endoderm; PAX6 for ectoderm) after directed differentiation in adherent cultures. Scale bar in **c**: 50 μm.

**Supplementary Figure 16.**
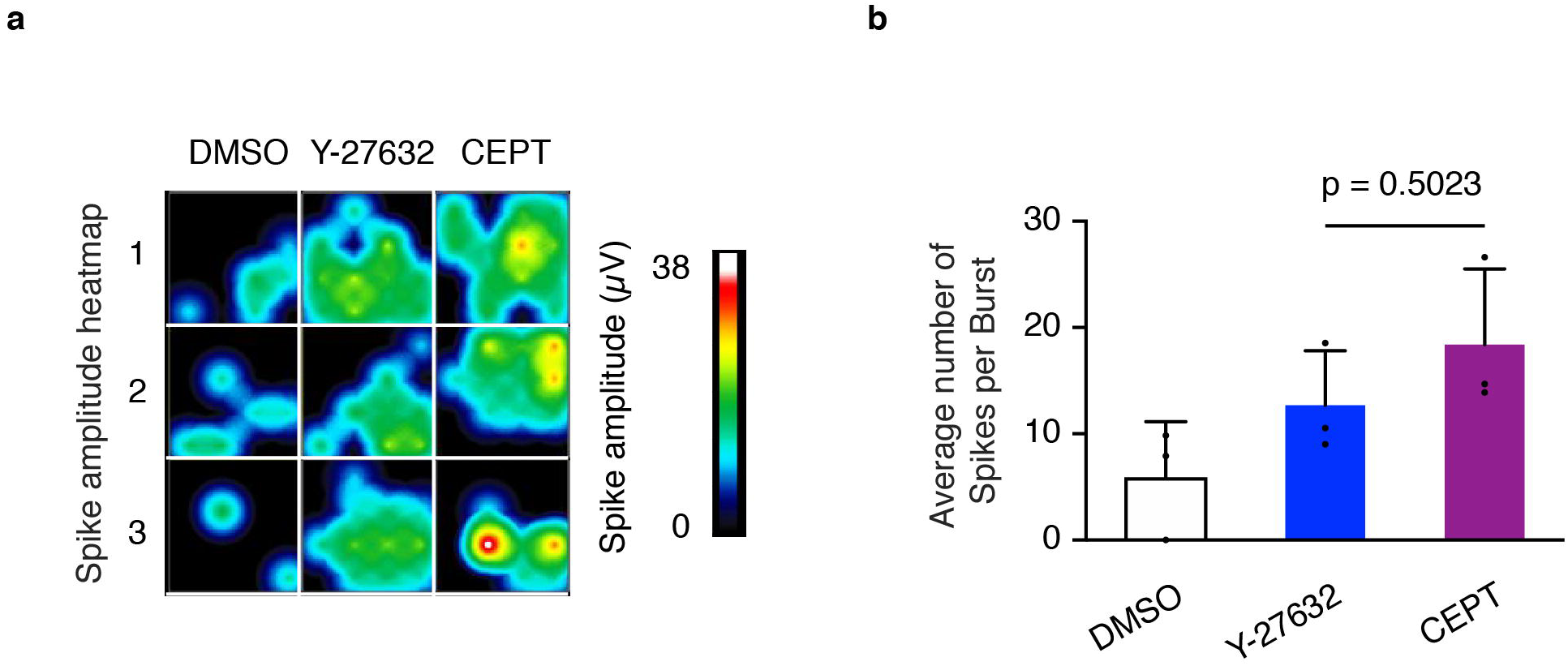
Improved motor neuron function after CEPT treatment. **a, b,** Frozen vials of iPSC-derived motor neurons were thawed and treated with Y-27632 and CEPT for 24 hr. Electrophysiological characterization of motor neurons was recorded using multi-electrode array technology (7 days post-thawing; n = 3 wells for each group). *P* = 0.5023, one-way ANOVA.

**Supplementary Figure 17.**
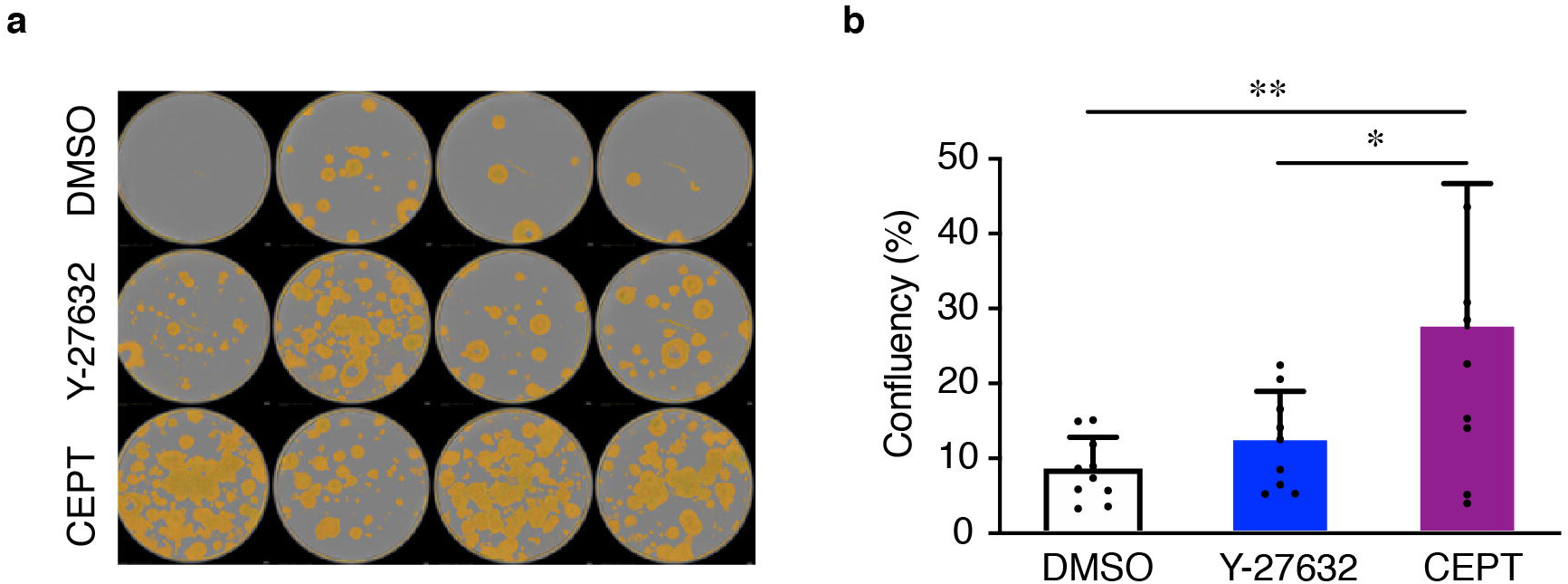
Improved iPSC line establishment by CEPT treatment. **a,** Human skin fibroblasts were reprogrammed using the Yamanaka factors and emerging individual iPSC colonies were manually picked and transferred to new plates. At day 8, cell confluency was measured demonstrating the superiority of CEPT for colony picking and establishment of new iPSC lines. **b,** Quantification of data shown in **a**. Data represent mean ± s.d. (n = 10 wells for each group), \*\**P* = 0.0044, \**P* = 0.0292, one-way ANOVA.

**Supplementary Figure 18.**
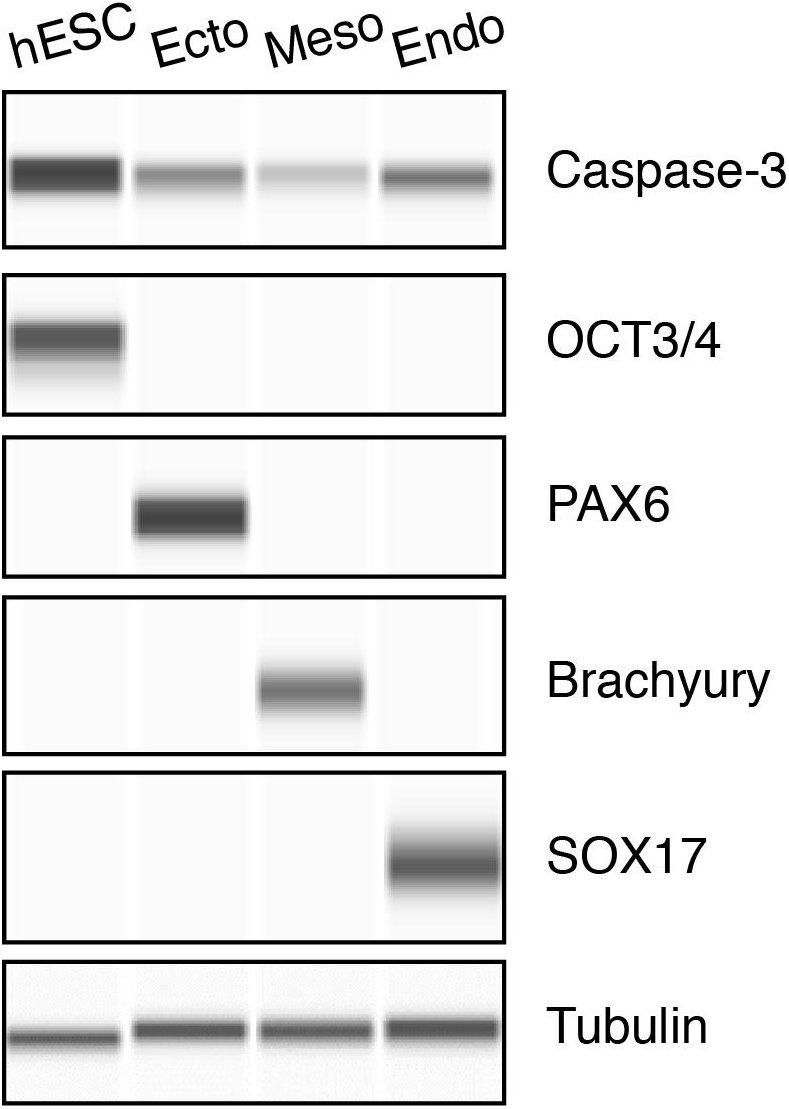
Caspase-3 expression in pluripotent and lineage-committed cells. Basal expression levels of Caspase-3 in hESCs (WA09) in comparison to their lineage-restricted precursors after directed differentiation into ectoderm (PAX6), mesoderm (Brachyury), and endoderm (SOX17). Note the strong Caspase-3 expression at the pluripotent state and downregulation upon differentiation.

**Supplementary Figure 19.**
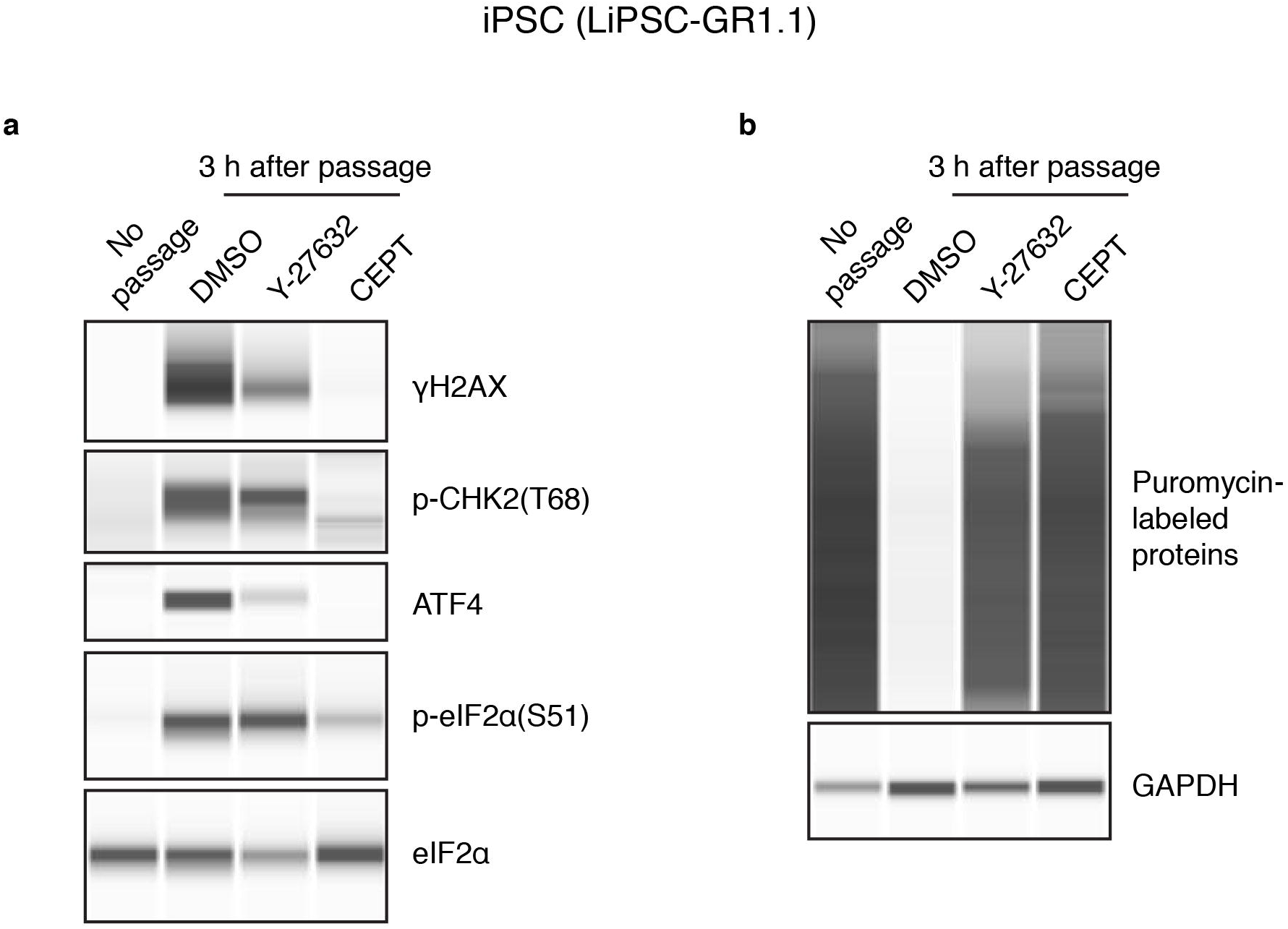
CEPT provides cytoprotection for iPSCs during cell passaging. **a,** Western blot analysis of iPSCs showing the dramatic changes that occurred at 3 h post-passage in the presence of DMSO and Y-27632. Note that CEPT mitigated these changes and cells were more similar to the condition prior to passage (“no passage” serving as control group). **b,** Puromycin pulse-chase experiment (50 min puromycin exposure) demonstrating that CEPT-treated cells showed higher protein synthesis capacity than cultures passaged with Y-27632. Note that protein synthesis appeared completely stalled in the presence of DMSO. All experiments were performed at 3 h post-passage.

**Supplementary Figure 20.**
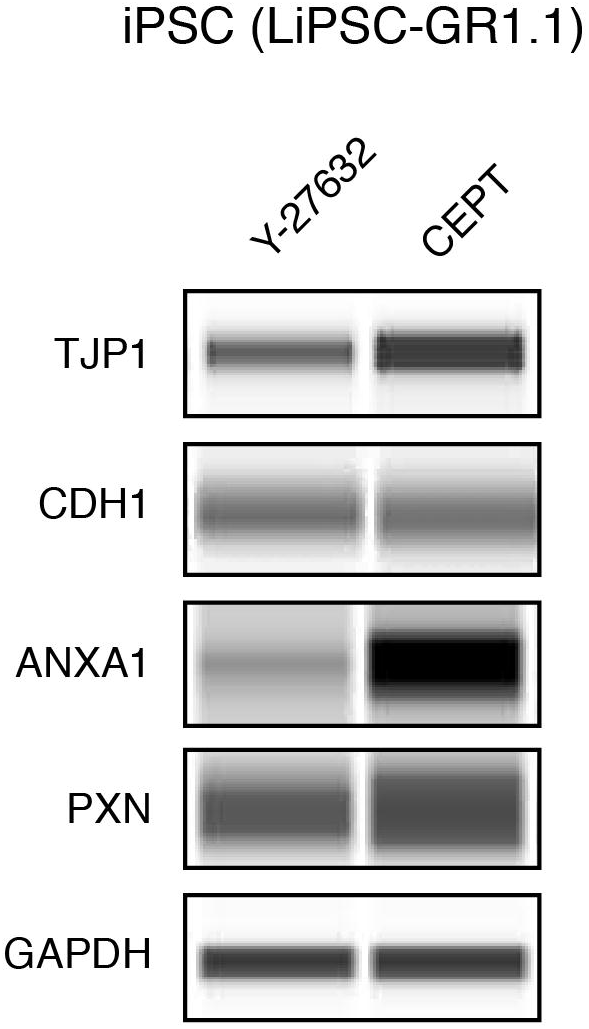
Western blot analysis cell membrane-associated proteins. Human iPSCs were dissociated and exposed to Y-27632 or CEPT for 24 h. All proteins are expressed at higher levels after CEPT treatment. GAPDH was used as a loading control. Similar results were obtained with hESCs (see Fig. 7f).

**Supplementary Figure 21.**
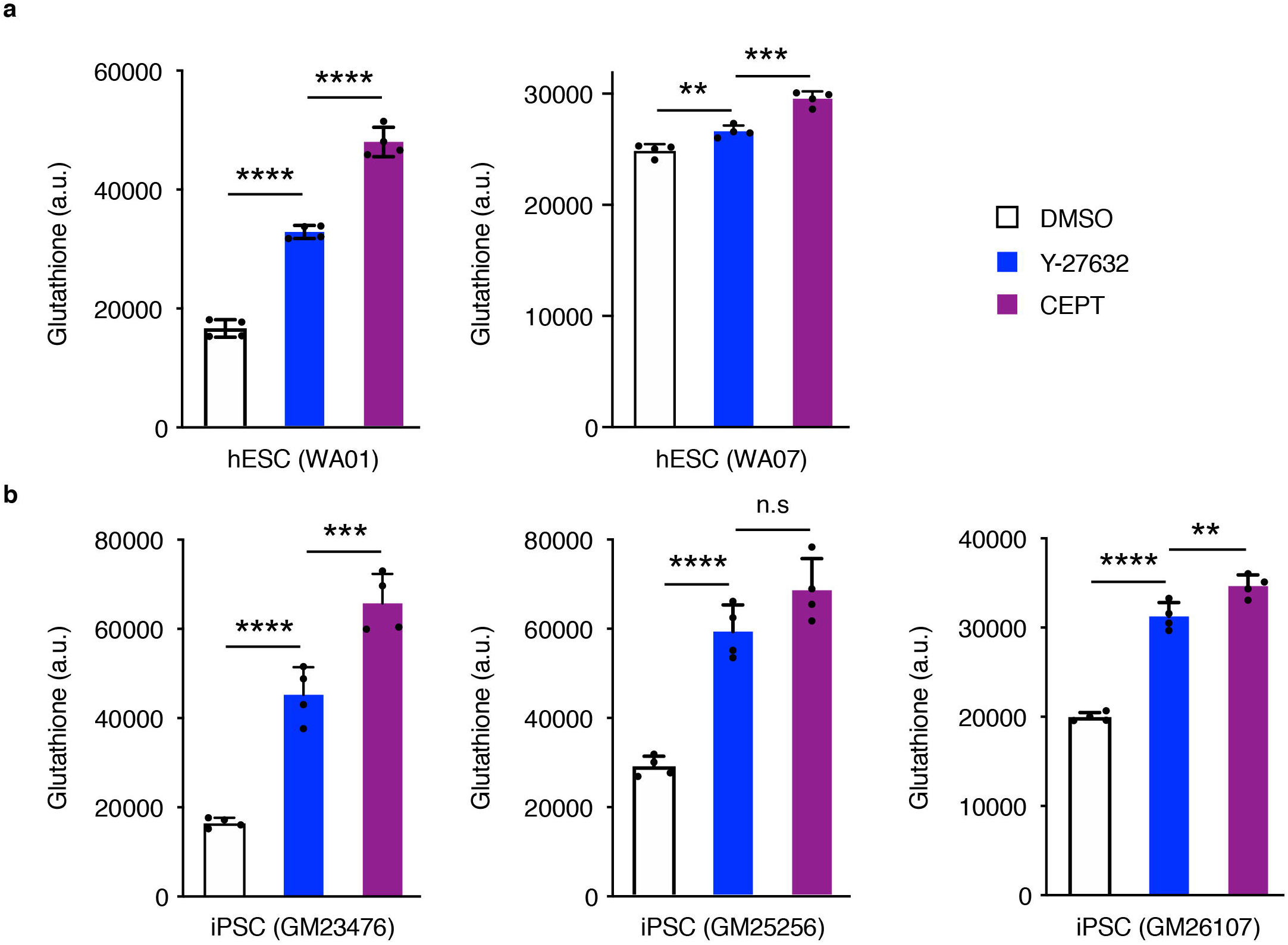
Measurement of glutathione levels in various hPSC lines. Glutathione levels were consistently higher in CEPT treated cultures compared to DMSO and Y-27632. Similar results were obtained for two hESC lines (**a**) and three iPSC lines (**b**) (n = 4 wells for each group). \*\*\*\**P* < 0.0001, \*\*\*\**P* <= 0.0010, \*\**P* <= 0.0075, one-way ANOVA.

**Supplementary Movie 1. Cell behavior during passaging is controlled by small molecules** Human ESCs (WA09) were dissociated with Accutase, plated on a vitronectin-coated 6-well plate in E8 medium, and treated with DMSO, Chroman 1, C+E, and Emricasan. Movies were recorded with a 10× objective at a 10 min interval. See the importance of Chroman 1 for cell survival and the inefficiency of Emricasan if administered alone. Scale bar, 25 μM.

### Supplementary Tables

**Supplementary Table 1. The 113 hits identified in the primary single agent screen.** Included for each hit are Sample Name, PubChem SID, CAS#, Dose-response curve classification, AC_50_, Efficacy, Max data, primary MOA and SMILES. Trans-ISRIB is appended to demonstrate its inefficiency to promote cell survival as a single agent.

**Supplementary Table 2. The 316 hits identified in ultra-low cell density screens.** Included for each hit are Sample Name, PubChem SID, CAS#, Dose-response curve classification, AC_50_, Efficacy, Max data, primary MOA and SMILES.

## Acknowledgements

We thank Paul Shinn, Misha Itkin, Zina Itkin, Carleen Klumpp-Thomas, John Braisted, Jessica Freilino, Ty Voss, Michael Iannotti, Charles Pepper Bonney, Yeliz Gedik, Deborah Ngan, Anna Rossoshek, and Ann Knebel for their support throughout this work. We are grateful to Alan Hoofring and Ethan Tyler from the NIH Medical Arts Design Section for their technical expertise. The authors would like to thank to David Panchision for critical reading of the manuscript.

## AUTHOR CONTRIBUTIONS

Y.C. and I.S. conceived the project. Experiments and Screening: Y.C., C.A.T., V.M.J., P.O., P-H.C., T.D., D.T., Y.F., S.M. Data Analysis and Discussions: Y.C., C.A.T., L.C., V.M.J., C.M., C.A.L., C.P.A., A.S., I.S. Manuscript writing: Y.C., A.S., I.S.

## Additional information

Extended results of small molecule screening are available here: https://ipsc.ncats.nih.gov/screening/

## Methods

### Cell culture

All human ESCs lines (WA01, WA07, WA09) and iPSCs (LiPSC-GR1.1^71^ and the following lines from Coriell Institute for Medical Research: GM25256^45^, GM23476, GM2610 and AICS-0023 clone 20) were maintained under feeder-free condition using Essential 8 (E8) medium and vitronectin (VTN-N) (Thermo Fisher Scientific). In some cloning experiments, StemFlex medium (Thermo Fisher Scientific) and laminin-521 (LN521; BioLamina) were used. Cells were routinely passaged using 0.5 mM EDTA diluted in phosphate buffered saline (PBS) without calcium or magnesium (Thermo Fisher Scientific) when cell culture plates reached ∼70-90% confluency, typically every 4 to 5 days. Some of the cell expansion experiments were performed using the automated CompacT SelecT cell culture system (Sartorius). Enzymatic cell dissociation was performed using Accutase (Thermo Fisher Scientific) and cell survival was tested by administration of different chemical compounds (see below) as indicated in respective experiments. For EB formation, hPSCs were single-cell dissociated using Accutase and plated into 6-well ultra-low attachment (ULA) plates (Corning) in Essential 6 (E6) medium (Thermo Fisher Scientific), 96-well ULA round bottom plates (Corning) and AggreWell plates (StemCell Technologies). For neural induction, hPSCs were treated with LDN-193189 (100 nM; Sigma-Aldrich) and A83-01 (2 μM; Tocris) in E6 medium for 6 days. Commercially available kits were used for mesoderm and endoderm induction (StemCell Technologies). All karyotyping experiments were performed by Cell Line Genetics (Madison, WI). All cultures were maintained at 37°C under humidified 5% CO_2_ and atmospheric O_2_. In some experiments (e.g. single-cell cloning), cell survival was also tested under hypoxic conditions (5% O_2_; data not shown). Human PSCs were cryopreserved in E8 medium, 10% DMSO, and either Y-27632 or CEPT. Mycoplasma testing was performed on a regular basis (every 2 weeks) using the Mycoalert Detection Kit (Lonza).

### Small molecule chemical compounds

The ROCK inhibitor Y-27632 (1254) and Trans-ISRIB (5284) were purchased from Tocris. Chroman 1 (HY-15392) was purchased from MedChem Express. Emricasan (S7775) was purchased from Selleckchem. Polyamine supplement (P8483) and antioxidant supplement (A1345) were purchased from Sigma-Aldrich. To prepare the CEPT cocktail the following concentrations were used: 50 nM Chroman 1, 5 μM Emricasan, Polyamine supplement diluted 1:1000 according to manufacturer’s recommendation, 0.7 μM Trans-ISRIB. The small molecule libraries used for HTS include the NCATS Pharmacologically Active Chemical Toolbox (NPACT) designed to inform on novel phenotypes, biological pathways and cellular processes; the NCATS Pharmaceutical Collection (NPC) including FDA-approved drugs; the Mechanism Interrogation PlatE (MIPE); and the commercially available libraries Tocriscreen Plus (Tocris) and LOPAC 1280 (Sigma-Aldrich). More information and details on NCATS libraries are available here: https://ncats.nih.gov/preclinical/core/compound

### Quantitative high-throughput screening (qHTS)

qHTS was performed on the Kalypsys robotic system (see also Supplementary Fig. 2). For screening in 1536-well format, plates with white flat bottom (Corning) were coated with VTN-N diluted in E8 media (2 μl per well) at 37 °C for 1 h and received 23 nl small molecule compounds in DMSO transferred from the compound library source plates by a pintool. Once 1536-well plates were ready, hESCs (WA09) were dissociated with Accutase, re-suspended in E8 medium and dispensed into the 1536-well plates at 500 cells/well (3 μl per well, resulting in a final assay volume of 5 μL in each well). After 24 h, 2 μl CTG was dispensed into each well and the luminescence signal was read using the ViewLux μHTS microplate imager (Perkin-Elmer). After normalizing all CTG values with that of 10 μM Y-27632 (positive control representing 100%; DMSO as negative control), we set a threshold of 20% for individually tested compounds and identified 113 primary hits (Supplementary Fig. 3a and Supplementary Table 1). These hits revealed a wide variety of dose–response curves that could be classified ^28^ into curve classes 1.1 (21 hits), 1.2 (4 hits), 2.1 (3 hits) and 2.2 (10 hits) (Supplementary Fig. 3b,c). When the cell viability assay was carried out at ultra-low cell density (10 cells/well) in 1536-well plates, hESCs were plated on LN521 and cultured in StemFlex medium to avoid daily media change. Dissociated hESCs were dispensed in the presence of Chroman 1 and Emricasan, and NPACT and kinase inhibitor libraries were tested at a single concentration of 10 μM in the primary screen. All plates were prepared in triplicates to compensate for cell dispensing variation. After ranking all compounds by their median CTG reading, 316 compounds were chosen for follow-up screens. In these follow-up screen, all compounds were tested at 7 different concentrations to generate dose-response curves and 6 replicates were used to compensate for cell dispensing variation.

### Combinatorial small molecule matrix screening

Cell viability assay was performed as described above (qHTS section), except that compound combinations were spotted using the Labcyte Echo 550 acoustic dispenser (Labcyte). Dose matrices were prepared in a 10 × 10 checkerboard formats, in which the two single agents were spotted in the last row and the last column, respectively (Example matrices are shown in Supplementary Fig. 7).

### HotSpot kinase inhibitor profiling

HotSpot kinase assays were performed by Reaction Biology (Malvern, PA) to determine IC_50_ for Y-27632 and Chroman 1 towards their primary targets including ROCK1 and ROCK2. Human kinase profiling against a panel of 369 human wild-type kinases was also carried out by Reaction Biology (Supplementary Figs. 5 and 6).

### Time-lapse video microscopy and fluorescence microscopy

Live-cell imaging was performed at 37°C under humidified 5% CO_2_ atmosphere using the IncuCyte Zoom system (Sartorius). Phase and fluorescence images were also acquired using Leica DiM8 and Zeiss LSM710 microscopes, as well as GE Healthcare IN CELL Analyzer 2200 high-content imaging system.

### Immunocytochemistry for cultured cells

Cell were fixed with 4% paraformaldehyde (PFA) for 30 minutes, permeabilized and blocked with 0.3% Triton X-100 and 5% BSA in PBS for 1 h. Rabbit anti-PAX6 antibody (BioLegend, 901301, 1:200), rabbit anti-brachyury antibody (Cell Signaling Technologies (CST), 81694, 1:200) and rabbit anti-SOX17 antibody (Abcam, 84990, 1:200) were used to stain ectoderm, mesoderm and endoderm progenitor cells. To stain cytoskeletal components, pluripotency and DNA damage markers, rabbit anti-MYH10 (Millipore-Sigma, ABT1340, 1:200), mouse anti-Oct3/4 human isoform A (BD Biosciences, 561628, 1:40) and rabbit anti-Phospho-Histone H2A.X (CST, 9718S, 1:250) were used. Primary antibody staining was followed by labeling with goat anti-mouse Alexa Fluor 488 (115-546-068, 1:400), donkey anti-rabbit Alexa Fluor 488 (711-545-152, 1:400), and donkey anti-rabbit Rhodamine (711-025-152, 1:600), goat anti-rabbit Alexa Fluor 647 (111-606-144, 1:400) conjugated secondary antibodies from Jackson ImmunoResearch Laboratories and Hoechst 33342 (Thermo Fisher Scientific, H3570, 1:2000). Actin was visualized by staining with Alexa Fluor 488-conjugated phalloidin (Thermo Fisher Scientific, A12379, 1:40, methanol stock solution prepared according to the manufacturer’s instructions).

### Live and dead cell assays

Simultaneous staining with CellTrace calcein green AM (Thermo Fisher Scientific, C34852, 1:2000) to indicate intracellular esterase activity and with red fluorescent propidium iodide (Thermo Fisher Scientific, P3566, 1:2000) to indicate loss of plasma membrane integrity allowed quick discrimination between live and dead cells. Caspase-3/7 activation in live cells was monitored using CellEvent Caspase-3/7 Green Detection Reagent (Thermo Fisher Scientific, C10423, 1:1000) in IncuCyte Live Cell Analysis System. Cells were plated at a density of 10^6^ cells/well in 6-well plates, treated with indicated compounds and imaged with a 10x objective. The CellTiter-Glo and CellTiter-Glo 3D assays (Promega, G9241, G9681) were carried out following the manufacturer’s instructions to quantify cell viability in adherent and EB cultures, respectively. Luminescence was read using the ViewLux μHTS microplate imager (Perkin Elmer).

### Western Blotting

The Wes automated Western blotting system (ProteinSimple) was used for quantitative analysis of protein expression and protein phosphorylation following the manufacturer’s instruction and all Western blot data are displayed by lanes in virtual-blot like images. Briefly, cell lysate and reagents were loaded into assay plates and placed into the Wes system. Cell lysates were loaded into the capillary automatically and separated by size as they migrate through a stacking and separation matrix. After the separated proteins were immobilized to the capillary wall, target proteins were identified using primary antibodies and immuno-probed using HRP-conjugated secondary antibodies and chemiluminescent substrate. The resulting chemiluminescent signal was detected by integrated WES detection camera and quantitated using the Compass software. Primary antibodies used are as follows: Caspase-3 (CST, 9665, 1:50), cleaved Caspase-3 (CST, 9664, 1:50), GAPDH (Santa Cruz, sc25778, 1:2000), GFP (Chromotek, 3H9, 1:50), lamin B1 (Abcam, 16048, 1:50), OCT4 (Santa Cruz, sc9081, 1:50), Brachyury (CST, 81694, 1:50), SOX17(Abcam, 84990, 1:50), PAX6 (BioLegend, 901301, 1:50), phospho-CHK2 (CST, 2197T, 1:50), *γ*H2AX (CST, 9718S, 1:50), phospho-eIF2A (Novus, NBP2-67353, 1:50), anti-puromycin (Millipore, MABE343, 1:50), tubulin (Novus, NB600-936SS, 1:500), tight junction protein 1 (Novus Biologicals, NBP1-85047, 1:20), E-Cadherin (Novus Biologicals, MAB1838, 1:100), Annexin A1 (R&D, MAP37701, 1:100), and Paxillin (Novus Biologicals, AF4259, 1:4).

### Protein translation assay

To monitor protein synthesis, we used a modified version of a previously described nonradioactive method^63^. Briefly, pluripotent cells were dissociated with Accutase for 10 min at 37°C. To ensure single-cell dissociation, the cell suspension was gently pipetted up and down 4 times and washed with PBS. Cells were then pelleted, resuspended, and seeded onto VTN-N-coated plates (25,000 cells/cm^2^) in E8 medium containing DMSO, Y-27632, or CEPT. Cells were allowed to adhere for 2 h and then pulsed with 1 μM puromycin for 1 h. Subsequently, cells were harvested using a cell scraper, pelleted, washed with PBS, flash-frozen and stored at −20°C. Cell pellets were lysed by sonication in RIPA lysis buffer (Thermo Fisher Scientific) supplemented with Halt protease inhibitor cocktail (Thermo Fisher Scientific) and centrifuged at 14,000g for 15 min. Protein translation was analyzed on the WES automated western blotting system (ProteinSimple) using anti-Puromycin antibody (1:50; Sigma-Aldrich, MABE343).

### Ultra-low cell density assay

hESCs (WA09) were single-cell dissociated with TrypLE (Thermo Fisher Scientific) and seeded into 6-well plates at a density of 25 cells/cm^2^. StemFlex medium was used to promote hPSC survival and colony formation. Compounds were applied for 72 hours. On day 6 after plating, hPSC colonies were stained with calcein green-AM and imaged using the INCell Analyzer 2200 with 2× objective to tile entire wells. Local threshold algorithm was applied to segment live cells from the background. hPSCs formed compact colonies on VTN-N allowed quantification of colony formation rate and colony size (Fig. 3e-g). We noticed that hPSCs were highly migratory on LN521 and therefore quantified the normalized cell confluency as cell survival index (Fig. 3d).

### Single-cell cloning using flow cytometry

hESCs (WA09) were single-cell dissociated with TrypLE, filtered through 40 μm cell strainer and sorted into VTN-N-coated 96-well plates using FACSAriaIII Fusion (BD Biosciences) with a 100-μm nozzle and FACSDiva software (BD Biosciences). Stringent gating strategy was implemented in both side scatter (SSC) and forward scatter (FSC) gates to exclude doublets. StemFlex medium was used to improve hPSC clonal formation by minimizing perturbation due to media change. Tested compounds were applied for the first 72 h following FACS sorting. On day 9 after plating, hPSC clones were stained with calcein green AM and imaged using the INCell Analyzer 2200 with 2× objective. Local threshold algorithm was applied to identify clones. When calculating single-cell clone formation rate in 96-well plates we factored in the 60% survival rate following FACS sorting by dividing the clone count in each plate by the number of 57, instead of 96. To further assess how the mechanical stress of the cell sorting process affects cell survival of hPSCs, we also tested the Sony SH800S cell sorter equipped with microfluidic sorting chips and noticed similar damage to hPSCs. In some single-cell cloning experiments the effect of 5% low oxygen at 37°C was tested in tri-gas CO_2_ incubators (data not shown).

### Clonal cell line generation following gene editing with CRISPR/Cas9

Gene editing with CRISPR/Cas9 was carried out following the protocol developed by the Allen Institute for Cell Science with some modifications. The parental iPSC line GM25256 was purchased from Coriell Institute for Medical Research. The donor plasmid to insert mEGFP into the N-terminus of lamin B1 was purchased from Addgene (Cat# 87422). TrueCut Cas9 protein v2 and TrueGuide 1-piece modified synthetic gRNA (GGGGTCGCAGTCGCCATGGC) were purchased from Thermo Fisher Scientific. iPSCs (GM25256) were dissociated into single cells using TrypLE and a cell pellet of 8 × 10^5^ cells was resuspended in 100 μl Neon Buffer R with 2 μg donor plasmid, 2 μg Cas9 protein, and sgRNA in a 1:1 M ratio to Cas9. Before addition to the cell suspension, the Cas9/sgRNA RNP was precomplexed for a minimum of 10 min at room temperature. Electroporation was with one pulse at 1300 V for 30 ms. Cells were then immediately plated onto LN521-coated 6-well dishes with StemFlex medium and CEPT was applied for the initial 24 h after plating. When transfected cells had recovered to ∼70% confluence (usually after 3–4 d), cells were harvested for FACS using TrypLE. The cell suspension (1.0 × 10^6^ cells/ml in StemFlex with CEPT) was filtered through a 40-μm mesh filter into polystyrene round-bottomed tubes. Cells were sorted using a FACSAriaIII Fusion with a 100-μm nozzle and FACSDiva software. Forward scatter and side scatter (height vs. width) were used to exclude doublets. Single cells were deposited into LN521-coated 96-well plates that were filled with CEPT-containing StemFlex medium. Half medium changes were performed with StemFlex medium (without CEPT) on day 3, day 6 and day 9 after FACS sorting. Most single-cell derived clones were picked on day 9 and transferred to 6-well plates for further analysis.

### Genetic screening with ddPCR

Genetic screening using ddPCR was performed following a previously published protocol^45^. Briefly, ddPCR was carried out using the Bio-Rad QX200 Droplet Reader, Droplet Generator, and QuantaSoft software (Bio-Rad). The reference assay for the two-copy, autosomal gene RPP30 was purchased from Bio-Rad (assay ID dHsaCP1000485, cat. no. 10031243). The hydrolysis probe–based PCR amplifications targeted to GFP (insert) and AMP (backbone) are as follows: GFP, primers (5′-GCCGACAAGCAGAAGAACG-3′, 5′-GGGTGTTCTGCTGGTAGTGG-3′) and hydrolysis probe (/56-FAM/AGATCCGCC/ZEN/ACAACATCGAGG/3IABkFQ/); AMP, primers (5′-TTTCCGTGTCGCCCTTATTCC-3′, 5′-ATGTAACCCACTCGTGCACCC-3′) and hydrolysis probe (/5HEX/TGGGTGAGC/ZEN/ AAAAACAGGAAGGC/3IABkFQ/). The GFP assay was run in duplex with the AMP assay as well as the genomic reference RPP30-HEX. The ratios of (GFP copies/μl)/(RPP30 copies/μl) were plotted against (AMP copies/μl)/(RPP30 copies/μl) to identify cohorts of clones for ongoing analysis.

### Genetic analysis with junctional PCR and Sanger sequencing

A single PCR reaction was used to amplify both the edited and wild-type allele of lamin B1 (forward primer: CTCGTCTTGCATTTTCCCGC, reverse primer: GACCGAGACCCTGTTCCTTC). PCR reactions were prepared using Q5 High-Fidelity 2X Master Mix (New England BioLabs, Ipswich, MA) and 10 ng genomic DNA in a final volume of 25 μl. Cycling conditions were as follows: 98°C for 30 s; (98°C for 10 s, 65°C for 30 s, 72°C for 60 s) × 40 cycles; 72°C for 10 min. PCR amplicons were analyzed by Sanger sequencing (Genewiz, NJ, USA) following agarose electrophoresis and gel purification.

### Whole Exome Sequencing

Pluripotent cells were collected by scraping and genomic DNA was extracted using the column-based DNeasy Blood and Tissue Kit (Qiagen) according to the manufacturer’s guidelines. DNA 260/280 ratios were assessed by Nanodrop. Whole exome sequencing (WES) comprised three passaging timepoints (P) for hESC (WA09) and LiPSC-GR1.1 cell lines independently for a total of six samples: hESC P26 (passage 26) without CEPT, hESC P36 with CEPT, hESC P46 with CEPT, iPSC P31 without CEPT, iPSC P41 with CEPT, and iPSC P51 with CEPT. Analysis was as follows and pipeline code, including details of QC filters, is also available at the aforementioned Github repository. The Burrows-Wheeler Aligner (BWA) aligned samples to the hg38 reference genome, samples were sorted with Samtools, and the Genome Analysis Toolkit (GATK) Best Practices were followed for germline short variant discovery (GATK version 3.5-1; Broad Institute; https://software.broadinstitute.org/gatk/best-practices). Annotation of SNPs and indels was done in ANNOVAR version 2018-04-16 for the following databases: hg38 RefSeq Gene, dbSNP150, 1000 Genomes Project August 2015 release (TGP), The Exome Aggregation Consortium (ExAC) version 0.3, Catalogue of Somatic Mutations in Cancer (COSMIC) 70, ljb26 (for SNPs only, containing whole-exome SIFT and PolyPhen2 HDIV), and ClinVar July 2015 release.

Annotation filtration and statistical testing was done in R. Variants were searched for damaging mosaic mutations^10^, cancer hotspots^37^, and TP53 specifically. The mosaic mutation positions were converted to hg38 positions with UCSC Genome Browser LiftOver tool. Pearson correlations were computed with 95% confidence intervals and Kruskal-Wallis rank sum tests were done on the variables: passage number (0, 10, 20), sample genotype (noncarrier, heterozygous carrier, or homozygous carrier), cell line (hESC or iPSC), TGP global population frequency, COSMIC presence or absence, and PolyPhen2 predicted effect. Rarity was defined as TGP <= 0.05 or unknown frequency. For indels, ClinVar predicted effect was included instead of PolyPhen2, which is unavailable for such variants. Correlation plotting was done in the R corrplot package.

### Fibroblast reprogramming and iPSC generation

Human fibroblasts from a healthy individual (GM00038) was purchased from the Coriell Institute for Medical Research. Reprogramming was performed using the CytoTune-iPS 2.0 Sendai reprogramming kit according to the manufacturer’s instructions (Thermo Fisher Scientific). Briefly, fibroblasts were plated at 20,000 cells/cm^2^ and cultured until they were 30-60% confluent. Cells were transduced with the Sendai reprogramming vectors at an MOI of 5:5:3 (KOS:hc-Myc:hKlf4). Cells were maintained in fibroblast media until passaged onto VTN-N-coated plates on day 7. On day 8, medium was changed to E8 medium and cells were cultured for additional 20 days. On day 28, colonies were picked using a 25-gauge needle to cut single colonies into 6-8 pieces and transferred onto VTN-N-coated plates containing E8 medium only or supplemented with Y-27632 or CEPT for 48 h. After 48 h, cells were kept in E8 medium only. Confluency was assessed 8 days after colony picking using the Celigo Imaging Cytometer (Nexcelom Biosciences).

### Viability analysis of cryopreserved hPSCs and differentiated cells

Commercially available iPSC-derived human cardiomyocytes (cat. R1017), hepatocytes (cat. R1027), and motor neurons (cat. R1051) were purchased from Fujifilm Cellular Dynamics International (Madison, WI, USA). Instructions of the manufacturer were followed for thawing, media preparation, coating of culture plates, and cell plating density. Cells were counted using Cellometer Auto 2000 (Nexcelom Biosciences) and plated using recommended media containing DMSO, Y-27632 or CEPT. Cell survival was measured by CTG and CellEvent Caspase-3/7 Green Detection Reagent was used to monitor apoptosis 24 h after thawing.

### Multi-electrode array (MEA)

Cardiomyocyte and motor neuron activity was analyzed using the Maestro multi-electrode array system (Axion Biosystems) according to the manufacturer’s protocol. Briefly, cardiomyocytes were thawed in a 37°C bead bath for 4 min, followed by addition of 9 ml of room temperature cardiomyocyte plating media containing DMSO, Y-27632 or CEPT and mixed gently. Cardiomyocytes were plated at a density of ∼63,000 cardiomyocytes/cm^2^. Forty-eight hours post-plating plating media was replaced with the appropriate volume of cardiomyocyte maintenance media without DMSO, Y-27632 or CEPT. Motor neurons were thawed in a 37°C bead bath for 2.5 min, followed by addition of 9 ml of room temperature complete maintenance media, pelleted and resuspended in room temperature complete maintenance media containing DMSO, Y-27632 or CEPT and mixed gently. Motor neurons were plated at a density of 5 million neurons/cm^2^ in complete media containing 10 μg/ml laminin. Twenty-four hours post-plating motor neuron maintenance media was replaced with maintenance media without DMSO, Y-27632 or CEPT and a 50% media exchange was performed every 2-3 days. Recordings were acquired at day 7 and 14 post-plating. For cardiomyocytes MEA plates were coated with 8 μl of 50 μg/ml fibronectin over the electrode recording area. For motor neuron experiments, MEA plates were coated with 0.1% polyethyleneimine (PEI).

### Single EB differentiation assay

Individual EBs were generated in 96-well ULA plates with 20,000 cells plated in each well and cultured in E6 medium for 7 days to allow for spontaneous differentiation. On day 7, mRNA from individual EBs was extracted using TurboCapture 96 mRNA kit (Qiagen), reverse-transcribed into cDNA using Sensiscript RT kit (Qiagen), and pre-amplified using TaqMan PreAmp Master Mix (Thermo Fisher Scientific) and Taqman probes for PAX6 (Hs01088114), Brachyury (Hs00610080) and SOX17 (Hs00751752), followed by qPCR quantification of PAX6, Brachyury, and SOX17. Cell number variation among EBs was normalized with the quantity of actin transcripts (Hs01060665).

### Generation of cerebral organoids

Whole-brain cerebral organoids were generated with a commercial kit (STEMdiff Cerebral Organoid Kit, STEMCELL Technologies, Catalogue Number 08570), which has been adopted and optimized based on previously published protocol^50^. In brief, iPSCs (LiPSC-GR1.1) were detached using Accutase (Thermo Fisher Scientific) and plated in 96-well ULA round bottom plates (Corning) at a density of 9000 cells per well in 100 μl of Embryoid Body Formation Medium (STEMdiff Cerebral organoid Kit, STEMCELL Technologies) with either addition of 10 μM Y-27632 (Tocris) or CEPT cocktail. Additional 100 μl of Embryoid Body Formation Medium was added every other day for 5 days. At day 5, or typically when embryoid bodies (EBs) reached between 400-600 µm in diameter with round and smooth edges, each embryoid body was transfer into each well of 24-well ultra-low attachment plate (Corning) in 0.5 ml of Neural Induction Medium (STEMdiff Cerebral organoid Kit, STEM CELL Technologies). 48 hours after transfer, or when neuroepithelium emerged, each organoid was embedded in 15 μl droplet of Matrigel (Corning) and transferred into 6-well ultra-low attachment plate (Corning) in Neural Expansion Medium (STEMdiff Cerebral organoid Kit, STEMCELL Technologies). After three days, the media was switched to Neural Maturation Medium (STEMdiff Cerebral organoid Kit, STEM CELL Technologies) and media was exchanged every 3 days.

### Histology and immunohistochemistry of cerebral organoids

Organoids were fixed in 4% PFA at room temperature for 20 min and washed with PBS three times before immersion in 30% sucrose at 4 °C overnight. Tissues were embedded in O.C.T. compound (Fisher Scientific), cut into 20-*µ*m sections, and mounted on microscope slides (Fisher Scientific) for staining. For histological analysis, sections were stained for hematoxylin/eosin followed by dehydration in ethanol and xylene and coverslipped in Permount mounting media (Fisher Scientific). For immunohistochemical analysis, sections were permeabilized and blocked with 0.3% Triton X-100 and 5% BSA in PBS for 1 h. Mouse anti-TUJ1 (BioLegend, 801202, 1:200), rabbit anti-SOX2 (Millipore-Sigma, AB5603, 1:100), and rabbit anti-FOXG1 (Abcam, AB18259, 1: 1000) were used, followed by labelling with Alexa-Fluor 488 (Thermo Fisher Scientific, A-21202, 1:500) and Alexa-Fluor 568 (Thermo Fisher Scientific, A10042, 1:500) conjugated secondary antibodies and coverslipped with ProLong™ Glass Antifade Mountant with NucBlue™ Stain (Thermo Fisher Scientific). Fluorescence images were taken with a Leica DMi8 microscope using appropriate filters.

### Statistics

Results are shown as mean ± s.d. Statistical tests included unpaired, two-tailed Student’s t-tests and one-way analysis of variance (ANOVA) for multiple comparisons. P values of 0.05 or less were considered to denote significance.

